# Chaperoning of the histone octamer by the acidic domain of DNA repair factor APLF

**DOI:** 10.1101/2021.12.24.474097

**Authors:** Ivan Corbeski, Xiaohu Guo, Bruna V. Eckhardt, Domenico Fasci, Wouter Wiegant, Melissa A. Graewert, Kees Vreeken, Hans Wienk, Dmitri I. Svergun, Albert J.R. Heck, Haico van Attikum, Rolf Boelens, Titia K. Sixma, Francesca Mattiroli, Hugo van Ingen

## Abstract

Nucleosome assembly requires the coordinated deposition of histone complexes H3-H4 and H2A-H2B to form a histone octamer on DNA. In the current paradigm, specific histone chaperones guide the deposition of first H3-H4 and then H2A-H2B(*1*–*5*). Here, we show that the acidic domain of DNA repair factor APLF (APLF^AD^) can assemble the histone octamer in a single step, and deposit it on DNA to form nucleosomes. The crystal structure of the APLF^AD^–histone octamer complex shows that APLF^AD^ tethers the histones in their nucleosomal conformation. Mutations of key aromatic anchor residues in APLF^AD^ affect chaperone activity in vitro and in cells. Together, we propose that chaperoning of the histone octamer is a mechanism for histone chaperone function at sites where chromatin is temporarily disrupted.

**one sentence summary:** Histone chaperone APLF assembles histones H2A-H2B/H3-H4 into histone octamers to deposit them onto DNA and form nucleosomes.

## Introduction

APLF (Aprataxin and Polynucleotide kinase Like Factor) is a DNA repair factor in non-homologous end joining (NHEJ) repair of DNA double-strand breaks(*6*–*8*), a critical pathway involved in immune responses and cancer biology(*9, 10*). APLF is recruited to DNA break sites through interactions with DNA-end binding protein Ku and the XRCC4-DNA Ligase IV complex to form a scaffold for the NHEJ machinery(*8, 11, 12*). In addition to its role as a scaffold, APLF has been shown to also have histone chaperone activity via its conserved C-terminal acidic domain (APLF^AD^) (*13*) (Fig. 1A and Fig. S1). The precise role of APLF as a histone chaperone during DNA damage repair is not fully understood. It has been suggested to play a role in recruitment and exchange of histone H2A variant macroH2A, but also to regulate the deposition of H3-H4 on DNA through specific binding of H3-H4 (*13*). We recently found that APLF^AD^ is intrinsically disordered and can bind H2A-H2B as well as H3-H4 with high affinity(*14*). Such promiscuous histone binding has been observed before for other histone chaperones(*15*–*19*) and argued to play a role in nucleosome assembly(*20*–*22*), but its structural basis and implications are not fully understood. In particular, these histone chaperones could challenge the notion that nucleosome assembly is a step-wise process in which first H3-H4 is deposited on the DNA and then H2A-H2B, with each step guided by specific histone chaperones (*1-5*). We therefore wanted to understand how APLF interacts with H2A-H2B and H3-H4 and determine its functional consequences.

**Fig. 1.**
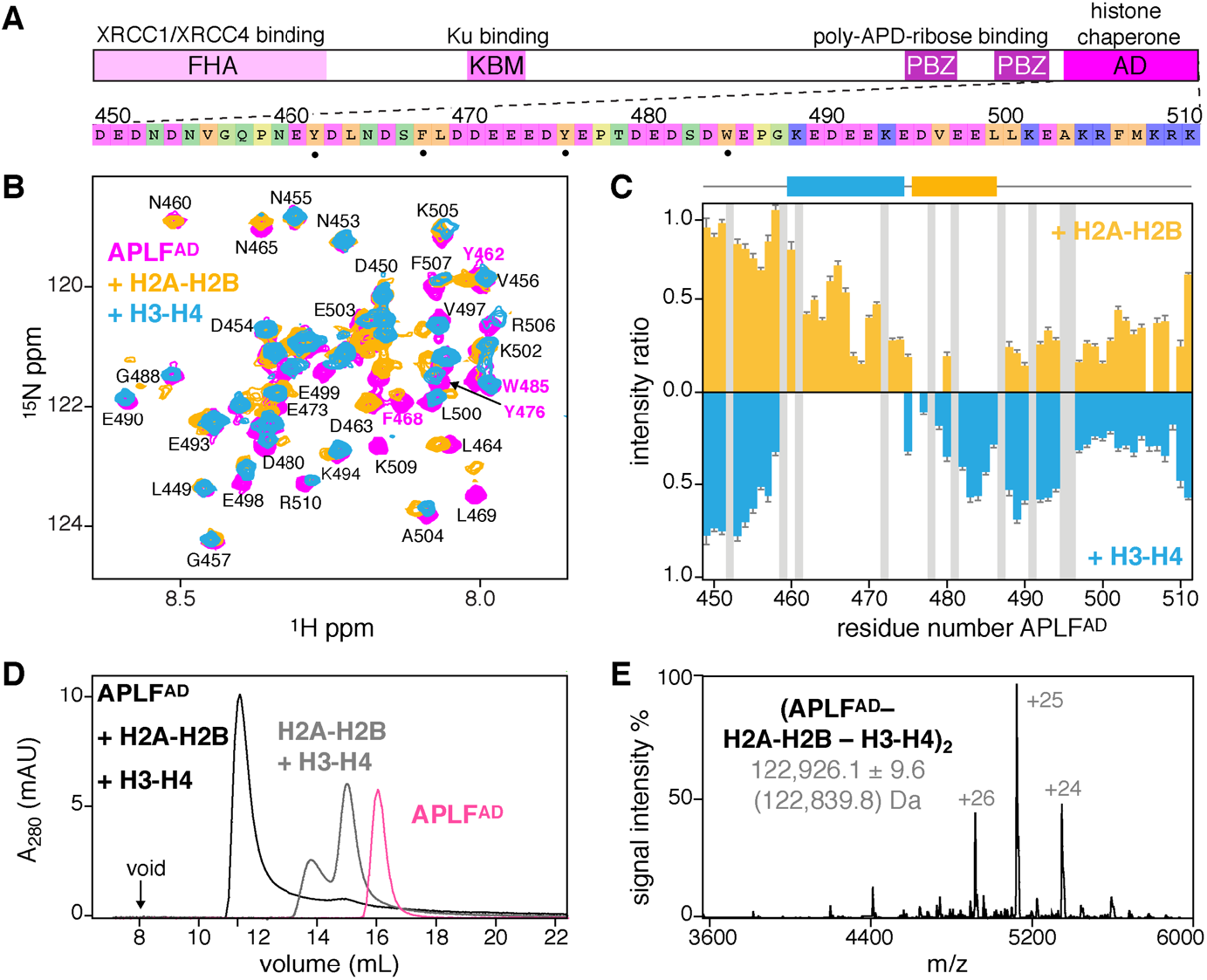
APLF^AD^ binds H2A-H2B and H3-H4 to form a (APLF^AD^–H2A-H2B–H3-H4)_2_ complex. **A**, APLF domain organization and domain function together with APLF^AD^ sequence, color coded according to amino acid properties. Key residues for histone interactions (•) indicated. **B**, Overlaid NMR spectra of APLF^AD^, free and bound to H2A-H2B or H3-H4. Aromatic residues labeled in magenta. Assignments from ref. (*14*). **C**, Relative APLF^AD^ NMR peak intensities upon binding H2A-H2B or H3-H4. Residues without data (prolines/overlapped peaks) in gray. The H2A-H2B (orange) and H3-H4 (blue) binding regions are indicated. **D**, SEC analysis of APLF^AD^ and octamer-mix (H2A-H2B + H3-H4) in absence and presence of saturating amounts of APLF^AD^. **E**, Native-MS spectrum of the APLF^AD^-histone complex with experimental (theoretical in brackets) molecular weights of the identified species.

## Results and discussion

### APLF^AD^ has distinct binding sites for H2A-H2B and H3-H4

We first wondered if H2A-H2B and H3-H4 bind to the same or to different sites in APLF^AD^. Using nuclear magnetic resonance (NMR) spectroscopy, we found that addition of either H2A-H2B or H3-H4 to APLF^AD^ resulted in severe peak intensity losses for distinct groups of residues, showing that APLF^AD^ contains two non-overlapping, adjacent binding regions for H3-H4 (residues N460-E474) and H2A-H2B (residues Y476-E486) (Fig. 1B,C). Both regions contain two aromatic residues. Previously we showed that Y476 and W485 are required for H2A-H2B binding(*14*), suggesting that Y462 and/or F468 could be crucial for H3-H4 binding. Other studies had however indicated W485 to be crucial for interaction with H3-H4 rather than H2A-H2B based on pull-down experiments at high salt(*13*). To resolve this, we performed isothermal titration calorimetry (ITC) experiments and found that Y462 is important for H3-H4 binding, while Y476A/W485A (named double anchor mutant DA-AB) had negligible influence on binding H3-H4 (Fig. S2). Further NMR experiments showed that H3-H4 binding involves the α1-α2 region of H3 (Fig. S2), analogous to our earlier results on H2A-H2B binding where binding entailed the α1-α2 region of H2A and H2B(*14*). These data indicate that APLF^AD^ has a distinct set of aromatic anchor residues to bind either H2A-H2B or H3-H4 in a specific manner.

### APLF^AD^ binds H2A-H2B and H3-H4 as a histone octamer complex

To test whether APLF^AD^ could bind to both H2A-H2B and H3-H4 simultaneously, we added APLF^AD^ to a stoichiometric mixture of H2A-H2B and H3-H4 (referred to as octamer-mix) and analyzed complex formation using size-exclusion chromatography (SEC) (Fig. 1D and Fig. S3). As expected, in the absence of other factors the mixture eluted as separate histone dimer and histone tetramer complexes. Strikingly, upon addition of APLF^AD^ a single high-molecular weight complex was obtained that contained all three components: APLF^AD^, H2A-H2B, and H3-H4 (Fig. S3). This indicated that two APLF^AD^ may be able to chaperone (i.e., bind) all histone components of the nucleosome at once. To further confirm this, we used native mass spectrometry (MS) and observed formation of a complex of 123 kDa, corresponding to two APLF^AD^ bound to two copies of H2A-H2B and H3-H4 each, (APLF^AD^–H2A-H2B–H3-H4)_2_ (Fig. 1E and Fig. S4). This complex has overall globular shape as shown from small-angle X-ray scattering experiments (Fig. S5) and is formed with sub-micromolar affinity (*K*_D_ ∼150 nM) as measured from ITC experiments (Fig. S6). Overall, these data demonstrate that APLF^AD^ can bind H2A-H2B and H3-H4 simultaneously to form a stable and high-affinity complex that contains the core histones at the same stoichiometry as found in the nucleosome.

### APLF^AD^ assembles H2A-H2B and H3-H4 as a native histone octamer

To understand how APLF chaperones core histones in the (APLF^AD^–H2A-H2B–H3-H4)_2_ complex, we set out to solve its structure. We obtained crystals of the complex reconstituted from tailless histones and a truncated APLF^AD^ construct corresponding to residues 449-490 (APLF^AD^). This truncation does not affect the binding affinity of APLF^AD^ for histones (see Fig. 4A below). We resolved the crystal structure of this complex at 2.35 Å resolution (Fig. 2, Fig. S7 and Table S1). The histones H2A-H2B and H3-H4 in the APLF^AD^-complex are arranged as in the nucleosome(*23, 24*) (0.56 Å backbone RMSD), involving histone-histone contacts across the H3-H3’ tetramerization interface, the H2B-H4 helical bundle, the H2A-H2A’ interface and the H2A docking domain to H3-H4 (Fig. S8). The structure has overall pseudo two-fold symmetry, where two APLF^AD^ flank the octamer, tethering H2A-H2B to H3-H4 within a histone half-octamer (Fig. 2A). APLF^AD^ imposes both a steric and electrostatic block at the binding sites of both nucleosomal DNA gyres (Fig. 2B,C).

**Fig. 2.**
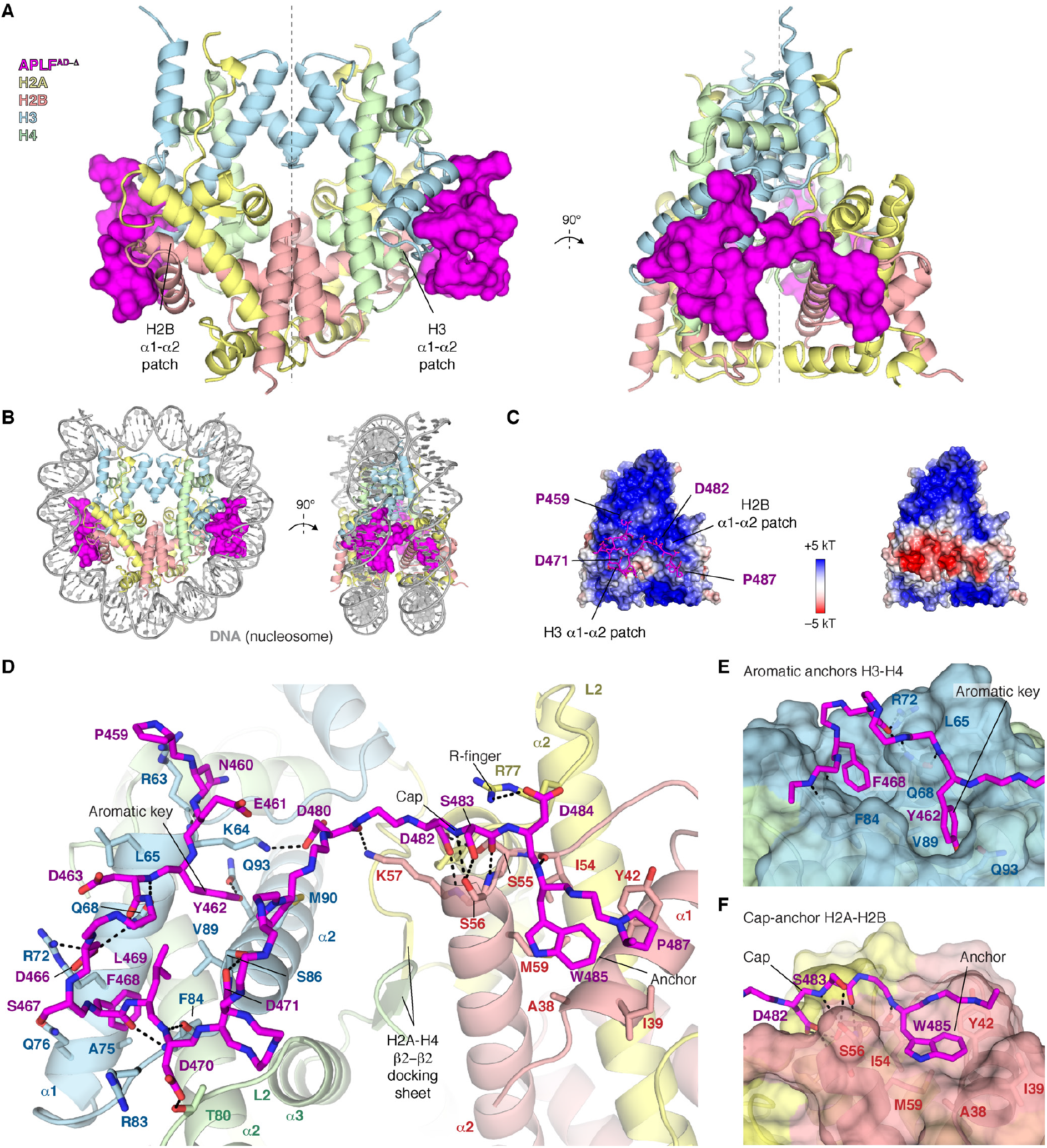
APLF^AD-Δ^ binds H2A-H2B and H3-H4 as a histone octamer. **A**, Ribbon view of the crystal structure of the APLF^AD-Δ^-histone octamer complex with APLF^AD-Δ^ (APLF^AD^ res. 449-490) shown as surface. Each APLF^AD-Δ^ binds primarily to the H2B and H3 α1-α2 patches (indicated) in each half-octamer. The pseudo-dyad axis is indicated with a dotted line. Color coding indicated in the Figure. **B**, Superposition of the APLF^AD^ –histone octamer complex and nucleosomal DNA highlighting the correspondence with the nucleosome structure and that APLF^AD^ blocks binding sites of both DNA gyres in the nucleosome. **C**, Electrostatic surface potential of the histone octamer (left) and the APLF^AD-Δ^–octamer complex (right), showing that APLF^AD-Δ^ binds the positively charged histone surface and creates a pronounced negatively charged bulk. **D**, Zoom on the APLF^AD^-histone octamer interface. Hydrogen bonds indicated as dashed lines; interface residues, the H2A-H4 docking β-sheet and known histone dimer binding motifs are labeled. **E**,**F** Zoom on the interaction of APLF^AD^ with H3-H4 (**E**) and H2A-H2B (**F**).

APLF^AD^ makes substantial interactions with both H2A-H2B and H3-H4, covering ∼800 Å^2^ of histone surface on H3-H4 and ∼400 Å^2^ on H2A-H2B (Fig. 2D). The regions involved in H3-H4 binding (residues P459-D471) and H2A-H2B binding (residues D482-P487) match well to the NMR results (Fig. 1C). The interaction is partially electrostatic, with an extensive network of intermolecular hydrogen bonds (Fig. 2D). Many of these interactions involve histone residues that otherwise bind the DNA phosphate backbone in the nucleosome, thus mimicking histone-DNA interactions (Fig. S8). In addition, aromatic residues of APLF^AD^ provide anchors that make extensive van der Waals interactions with the histones. APLF residues Y462 and F468 protrude deeply into hydrophobic pockets on the H3 α1-α2-patch (Fig. 2D,E), consistent with their role in H3-H4 binding (Fig. S2). Similarly, APLF W485 anchors to a shallow hydrophobic pocket on the H2B α1-α2-patch (Fig. 2D,F), in line with its role in H2A-H2B binding(*14*). Electron density for residues (E472-E481) that connect the H2A-H2B and H3-H4 binding regions is incomplete or missing in all but one of the APLF^AD^ chains (Fig. S7), suggesting that this segment forms a flexible linker. Overall, APLF^AD^ makes use of multiple known histone dimer binding modes: the cap-anchor(*25*) and the R-finger interaction(*26*) for H2A-H2B, and the aromatic-key motif(*27*) for H3-H4 binding (Fig. 2E,F). Structural comparison to other histone chaperones (Fig. S9 and S10) shows that APLF^AD^ uniquely combines these binding modes to bind H2A-H2B and H3-H4 simultaneously in their native histone octamer configuration.

### APLF^AD^ envelops the histone octamer with its C-terminal region

We next examined the binding mode of the full APLF^AD^ to full-length histones in solution using NMR and MS. NMR titration experiments of the octamer-mix to APLF^AD^ resulted in drastic peak intensity losses for all APLF^AD^ residues, except the N-terminal ten residues (Fig. 3A and Fig. S11). This suggests that the C-terminal residues 488-511, which are missing in the crystal structure, are also involved in binding, while the N-terminal region remains highly flexible in the complex. As the bound APLF^AD^ was not observable by NMR directly, we probed its conformation using relaxation dispersion experiments, exploiting the continuous interconversion of free and bound states. These experiments indicated that residues involved in H2A-H2B and H3-H4 binding experience significant changes in their chemical environment between free and bound states (Fig. 3B), supporting the crystal structure binding mode. Moreover, the fitted chemical shift differences indicate that residues in the C-terminal region are similarly engaged in binding and undergo a concerted binding event with the H2A-H2B and H3-H4 binding motifs (Fig. 3B and Fig. S11). This is further supported by cross-linking MS (XL-MS) experiments revealing reproducible cross-links between lysine residues within the APLF^AD^ C-terminal region and the H2B αC-helix (Fig. 3C). By including the NMR and XL-MS data, we extended the APLF^AD-^-histone octamer crystal structure into a model of the full-length APLF^AD^ bound to the histone octamer. The residues missing in the crystal structure were built in random coil conformation while imposing a maximum 27 Å Cα-Cα distance for the cross-linked residues in the APLF^AD^ C-terminal region and the H2B αC-helix. These restraints are compatible with a wide range of conformations of the C-terminal region in the final model, all in close proximity to the H2B αC-helix (Fig. 3D and Fig. S11). APLF^AD^ thus envelops the histone octamer completely, except for the central region around the dyad. In this binding mode, APLF^AD^ could influence the DNA interactions of the histone octamer and favor DNA binding at the dyad, the central region in the nucleosome.

**Fig. 3.**
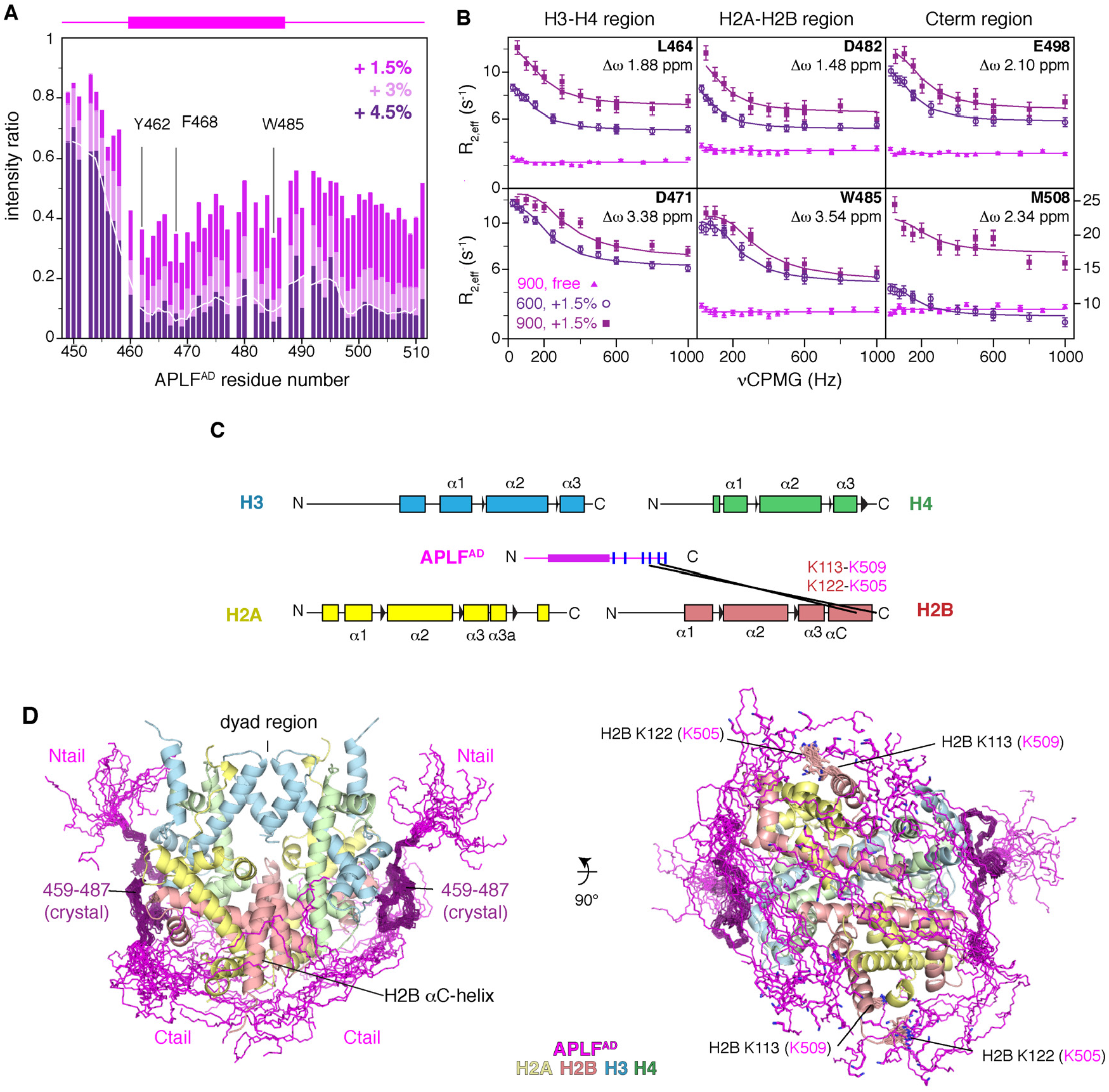
APLF^AD^ including the N- and C-terminal tails envelops the histone octamer. **A**, Relative NMR peak intensities of APLF^AD^ upon addition of octamer-mix show similar signal intensity decrease for the C-terminal region (residues 488-511) as for the histone binding region, indicating it is bound to the octamer surface. The 3-residue moving average intensity for the +4.5% data is shown as a white line. The APLF^AD^ fragment visible in the crystal structure is shown as a purple box on top of the Figure. Selected residues are labeled. **B**, Residues in the C-terminal region experience significant changes in their chemical environment and undergo a concerted binding event with the H2A-H2B and H3-H4 binding motifs, based on fitting of NMR relaxation dispersion data (see Fig. S11). **C**, The APLF^AD^ C-terminal region is in proximity of H2B as based on analysis of intermolecular lysine cross-links (black lines) in the APLF^AD^-histone octamer complex identified by mass spectrometry. Secondary structures of the histones are indicated. The APLF^AD^ fragment visible in the crystal structure is shown as a purple box. Lysine residues within APLF^AD^ are indicated as blue lines. **D**, Superposition of the twenty best ranking models of the (APLF^AD^–H2A-H2B–H3-H4)^2^ complex, showing that APLF^AD^ covers most of the DNA binding surface on the histone octamer with exception of the dyad region. The crystallized part of APLF^AD^ is shown in dark purple, the APLF^AD^ N- and C-terminal tails in magenta are modeled based on the intermolecular cross-links (cross-linked H2B residues shown as sticks and labeled in the right panel, APLF^AD^ residues in brackets).

### APLF^AD^ aromatic anchors are critical for histone octamer binding in vitro and in cells

To test the importance of the H2A-H2B and H3-H4 interactions of APLF^AD^ for binding and chaperoning of the histone octamer, we mutated the aromatic anchor residues that are involved in binding based on the octamer complex structure or implicated in binding isolated H2A-H2B(*14*) or H3-H4 (Fig. S2). We used double anchor mutants to disrupt either the H3-H4 interface (Y462A/F468A, DA-34) or the H2A-H2B interface (Y476A/W485A, DA-AB), and a quadruple anchor (QA) mutant that combines these mutations (see Table S3 for an overview of the mutants). Indeed, these mutations reduced the binding affinity of APLF^AD^ to the octamer-mix up to five-fold (Fig. 4A). Additionally, removal of the anchor residues resulted in a large decrease of binding enthalpy, suggesting a reduction in buried surface and thus a defect in assembly of the histone octamer (Fig. S12). Though deletion of the C-terminal region (Δ) did not affect binding affinity to the octamer-mix, reduced binding affinity and enthalpy was observed when the deletion was combined with the QA mutant (QA-Δ) (Fig. 4A) These data further indicate that the C-terminal region is involved in weak, dynamic interactions with the histone surface. To further probe the importance of the aromatic anchor residues for chaperone activity we tested wild-type and mutant APLF^AD^ in their ability to prevent non-native histone-histone contacts using XL-MS. As expected, for the octamer-mix alone, most histone-histone cross-links obtained are incompatible with the histone octamer structure (Fig. 4B). Strikingly, addition of wild-type APLF^AD^, but not aromatic anchor mutants, dramatically reduced the number of incompatible cross-links, further substantiating that APLF^AD^ functions as a histone chaperone and stabilizes the core histones in their nucleosomal octameric arrangement (Fig. 4B).

**Fig. 4.**
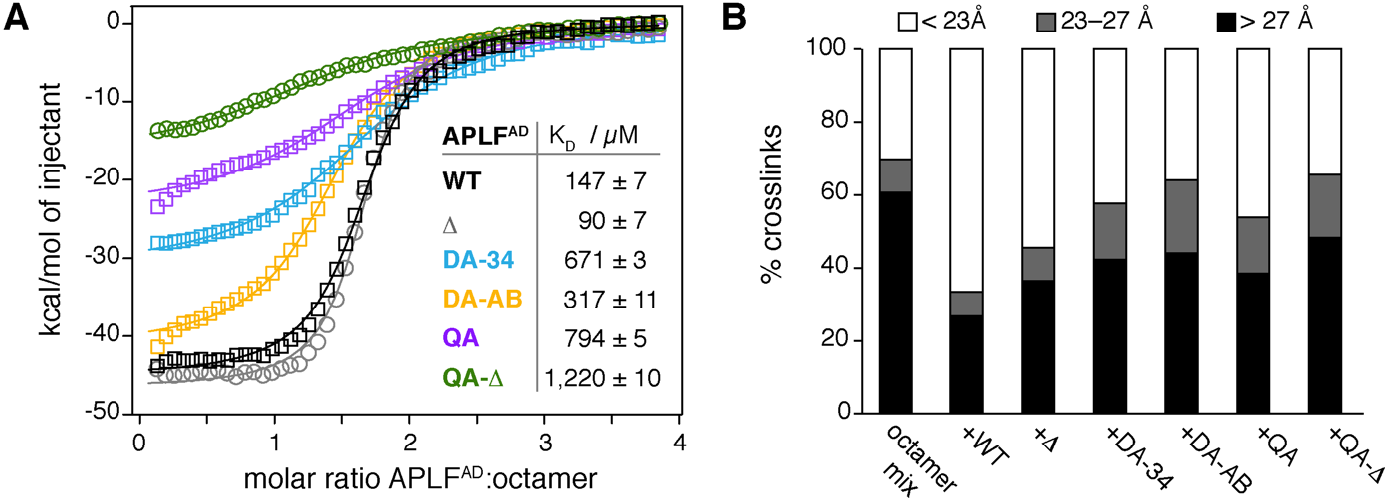
APLF^AD^ aromatic anchor residues are required for histone octamer assembly and chaperone activity in vitro. **A**, ITC binding curves and derived affinities (*K*^D^) of APLF^AD^ wild-type (WT), the truncation mutant used for crystallization (APLF^AD-Δ^, Δ) or the double (Y462A/F468A = DA-34; Y476A/W485A = DA-AB) and quadruple (Y462A/F468A/Y476A/W485A = QA) mutants titrated to octamer-mix. **B**, Percentage of octamer-compatible and -incompatible histone-histone lysine cross-links based on surface accessible Cα-Cα distances in the nucleosomal structure (PDB:2PYO) identified by XL-MS.

Our data show that the APLF acidic domain binds the histone octamer and we identify specific aromatic residues crucial for this interaction. We therefore wondered if this binding occurs in cells and how it affects APLF function during DNA damage. Consistent with previous results(*13*), we found that histone binding in cells is dependent on the presence of the acidic domain, using immunoprecipitation pulldown assays in the presence of benzonase (Fig. 5A,B). Mutation of the aromatic anchors in the DA-34 and DA-AB mutants result in both reduced H3-H4 and H2A-H2B binding. Combined mutation of all anchors in the QA construct was sufficient to fully abrogate all histone binding, indicating that the aromatic anchor interactions as captured in the crystal structure are crucial for APLF’s histone binding in cells. These data indicate that APLF may engage histone octamers rather than separate H3-H4 or H2A-H2B units, and they also suggest that H3-H4 and H2A-H2B binding may be interlinked in cells, consistent with the handling of octamers.

**Fig. 5.**
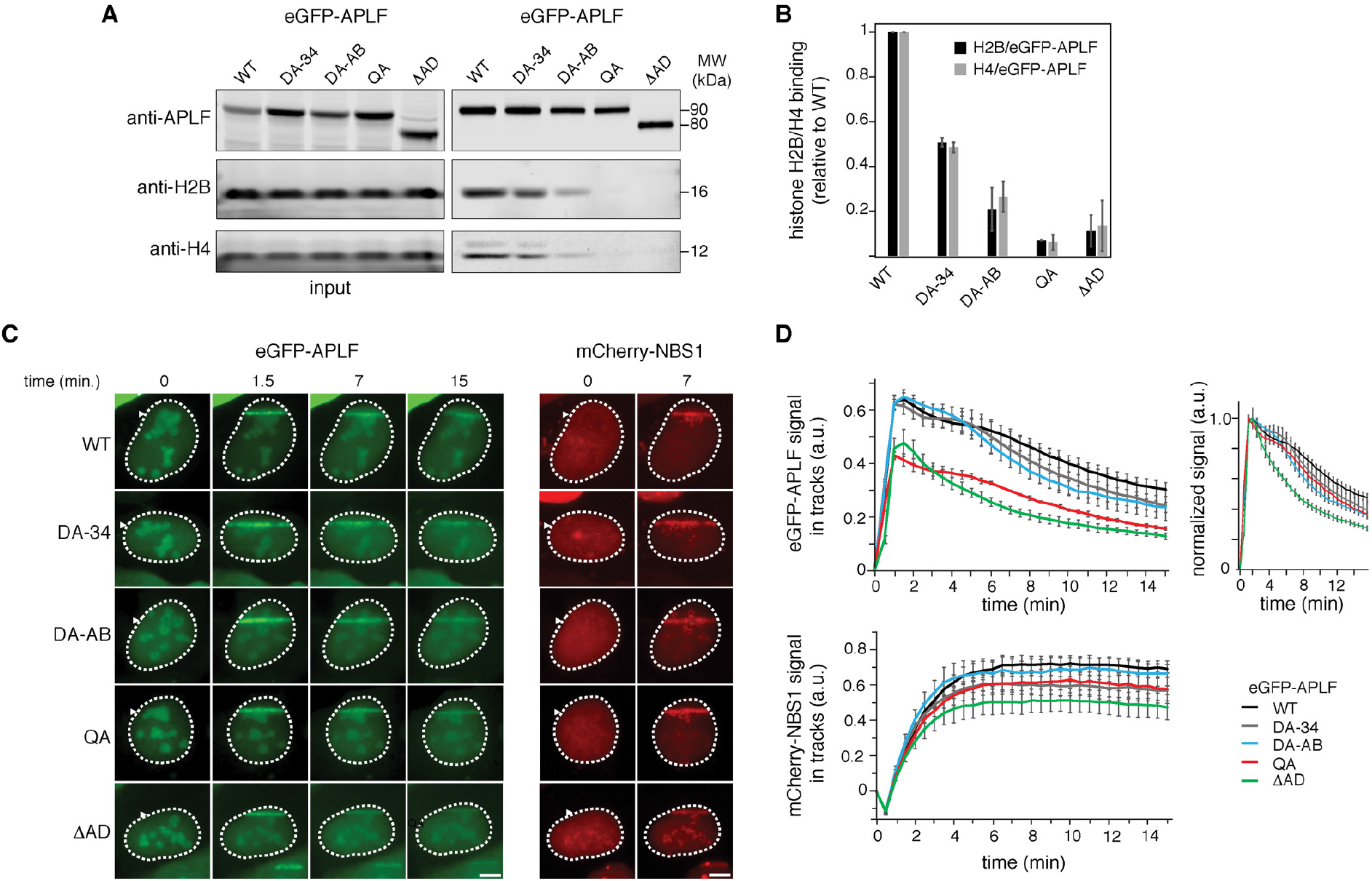
APLF^AD^ aromatic anchor residues are required for histone binding and recruitment to DNA damage sites in cells. **A**, Pull-downs of eGFP-APLF wild-type (WT), DA-34 (Y462A/F468A), DA-AB (Y476A/W485A), QA (Y462A/F468A/Y476A/W485A), or Δ AD (acidic domain deletion) mutant in the presence of benzonase following transient expression in U2OS cells. Blots were probed for GFP, H2B and H4. **B**, Average normalized H2B or H4 signal (with standard deviation) of duplicate pull-down results. Signals of H2B or H4 were normalized to that for eGFP-APLF WT or mutant protein level, then normalized to that of WT, which was set to 1. **C**, Live-cell imaging of the recruitment of eGFP-APLF WT, DA-34, DA-AB, QA or Δ AD mutant to DNA damage tracks generated by UV-A laser micro-irradiation in BrdU-sensitized U2OS cells (left panel). mCherry-NBS1 was co-transfected with eGFP-APLF. Live-cell imaging of the recruitment of mCherry-NBS1 in these cells is shown (right panel). Representative images are shown. Scale bars: 5μm. **D**, Quantification of the recruitment of eGFP-APLF WT or mutant protein (top left panel), and mCherry-NBS1 (bottom panel) to DNA damage tracks in cells from **C**. Normalized data in top right panel highlights relative differences in release kinetics. Data represent the mean values ± standard error of the mean (SEM) from 50 WT-or 30 mutant eGFP-APLF-expressing cells acquired in 3 independent experiments.

Previous work also showed that deletion of the acidic domain interferes with recruitment of APLF at DNA damage sites(*13*). Under conditions where accumulation of the DNA double-strand break marker NBS1 is clearly visible, we found that the acidic domain deletion mutant shows strongly reduced accumulation level at DNA damage sites (Fig. 5C,D). This effect is retained in the QA mutant, in agreement with the crucial role of the aromatic anchors for the function of the acidic domain in histone binding (Fig. 5A,B). The DA-AB and DA-34 mutants, which have only impaired histone octamer binding (see Fig 5A,B), showed no change in recruitment level. This suggests that their residual histone binding is sufficient to support accumulation at DNA damage sites. Together, these results indicate that APLF’s ability to bind the histone octamer is critical for its recruitment and retention at sites of DNA damage, thereby likely impacting DNA damage repair.

### APLF^AD^ chaperones the histone octamer to promote nucleosome assembly

Having established that APLF binds the histone octamer in vitro and in cells, we investigated whether APLF can deposit octamers on DNA to form nucleosomes, possibly to restore chromatin after DNA damage repair. We first tested if APLF^AD^ prevents non-native histone-DNA contacts, as expected for a histone chaperone(*1*), using a precipitation-rescue assay(*14, 28*). Indeed, APLF^AD^ rescued the precipitation of histones on DNA in a way that depends on the presence of the key aromatic anchor residues (Fig. 6A). This demonstrates that APLF^AD^ functions as a *bona fide* histone chaperone using the binding mode observed in the crystal structure. Next, we tested whether APLF^AD^ facilitates nucleosome formation. Using the nucleosome assembly and quantitation (NAQ) assay(*29*), we monitored nucleosome formation upon incubation of the octamer-mix with 207 bp DNA fragments in the presence of APLF^AD^, followed by digestion with micrococcal nuclease (MNase). Addition of APLF^AD^ to the histones caused increased protection of DNA fragments of 125-160 bp in a dose-dependent manner, consistent with nucleosome formation (Fig. 6B,D and Fig. S13). Together, these data indicate that APLF^AD^ prevents spurious histone-DNA interactions and allows deposition of the histone octamer on DNA to form nucleosomes.

**Fig. 6.**
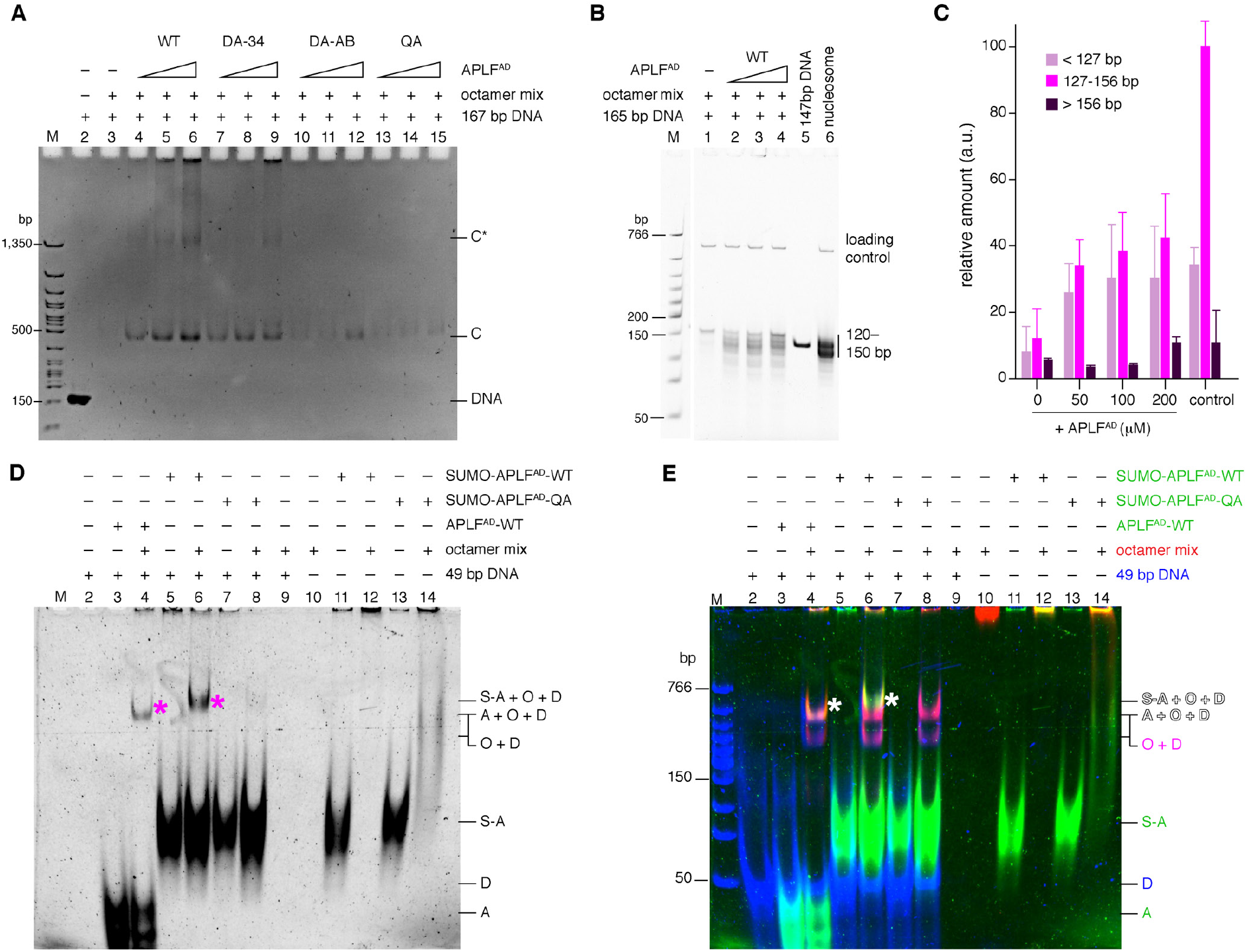
APLF^AD^ chaperones the histone octamer to promote nucleosome assembly. **A**, Native PAGE analysis of precipitation-rescue assay showing formation of soluble protein-DNA complexes (bands ‘C’ and ‘C*’) upon addition of increasing amounts of WT APLF^AD^ to octamer-mix with DNA, which is strongly reduced for mutant APLF^AD^. Band ‘C’ corresponds to the electrophoretic mobility of nucleosomes (Fig. S13). Total DNA control shown in lane 2. **B**,**C** NAQ results showing MNase digestion products obtained for octamer-mix with DNA and increasing amounts of APLF^AD^. Quantification of DNA digestion products (average and standard error of the mean (*n*=3)) in panel **C**. APLF^AD^ increases the protected nucleosomal bands (127-156 bp) (*p* = 0.018, 0.018, 0.016 for 50, 100, 200 µM) according to a one-tailed Students’ *t*-test. Salt-assembled nucleosomes are used as control. **D**,**E** Native PAGE analysis of indicated mixtures of APLF^AD^, octamer-mix and DNA, crosslinked with DSS. Panel **D** shows the Cy3-scan with APLF^AD^ signal before DNA staining, panel **E** shows a merged image of the APLF^AD^ (green), histone (red) and DNA (blue) scans (see Fig. S14 for individual scans). APLF^AD^ forms a ternary complex with histones and DNA (asterisk in lane 3 and 6). SUMO-APLF^AD^ was used as control to shift the ternary band above the background octamer-DNA complex. A = APLF^AD^, S-A = SUMO-APLF^AD^, O = histone octamer, D = DNA. Labels in **E** are color-coded according to the fluorescent dye.

We next sought to understand how APLF^AD^ may deposit octamers on DNA. As APLF^AD^ does not cover the entire DNA binding surface of the histone octamer (see model in Fig. 3), we hypothesized that the APLF^AD^-octamer complex would be able to bind a short piece of DNA, forming a ternary APLF^AD^-octamer-DNA complex representing a reaction intermediate during octamer deposition. Using fluorescently labeled proteins and cross-linking to trap transient complexes, we confirmed that APLF^AD^ alone does not bind DNA (Fig. 6D,E lane 2 vs. 3), while addition of a 49-bp DNA fragment to the histone octamer-mix alone resulted in precipitation (Fig. 6D,E lane 9). Upon incubation of the APLF^AD^-histone octamer complex with the DNA fragment, we detected a ternary complex containing DNA, histone octamer, and APLF^AD^ (Fig. 6D,E marked band in lane 4 and 6 and Fig. S14). Importantly, this ternary complex is not formed when using the APLF^AD^ QA mutant (Fig. 6D,E lane 8), indicating that in absence of proper APLF^AD^-histone binding the intermediate cannot be formed. Moreover, in the presence of a longer 147-bp DNA fragment the ternary complex could not be detected, but only DNA-histone complexes (Fig. S15), in line with the octamer being deposited on this longer DNA (as in Fig. 6B) and APLF^AD^ leaving the nucleosome product. Therefore, the APLF^AD^-octamer-DNA complex isolated with a short DNA fragment may represent an intermediate in the octamer deposition process by APLF.

Together, these data lead to a compelling model for nucleosome formation by APLF^AD^ that contrasts sharply with the stepwise nucleosome assembly pathway used by the other ATP-independent histone chaperones characterized so far(*30*) (Fig. 7). Our data showed that APLF^AD^, as a flexible and disordered protein, can bind both H2A-H2B and H3-H4 simultaneously, tethering them into a histone octamer in its nucleosomal configuration. In the complex, APLF^AD^ stabilizes the histone octamer and prevents non-native histone-DNA interactions. We propose that the exposed histone dyad region, where the octamer has highest affinity for DNA(*31*) and where DNA binding stabilizes the H3-H3’ interface of the (H3-H4)_2_ tetramer, allows the interaction with DNA to initiate octamer deposition. The DNA may then displace APLF^AD^ as it wraps around the histone octamer to form the nucleosome (Fig. 7). Notably, the APLF acidic domain is flexible and exposed within the core NHEJ complex(*11*), which can contain two copies of APLF through binding the DNA-end binding protein Ku80(*12*). The acidic domain of APLF could thus provide the NHEJ machinery with the capacity to assemble or capture histone octamers, store them during the DNA repair process, and to promote nucleosome assembly to restore chromatin after repair. As APLF binds conserved surfaces on the histones, it can likely also assemble histone octamers containing histone variants, consistent with identification of H2A.X and macroH2A in co-immuno-precipitation experiments with APLF(*13*). Furthermore, as APLF can bind PARylated histones through its PBZ domains, PAR binding could assist APLF’s ability to bind histones and stimulate APLF’s histone octamer chaperone activity. It is tempting to speculate that APLF could capture PARylated histones that are evicted from chromatin, and then assemble these via its acidic domain into histone octamers.

**Fig. 7.**
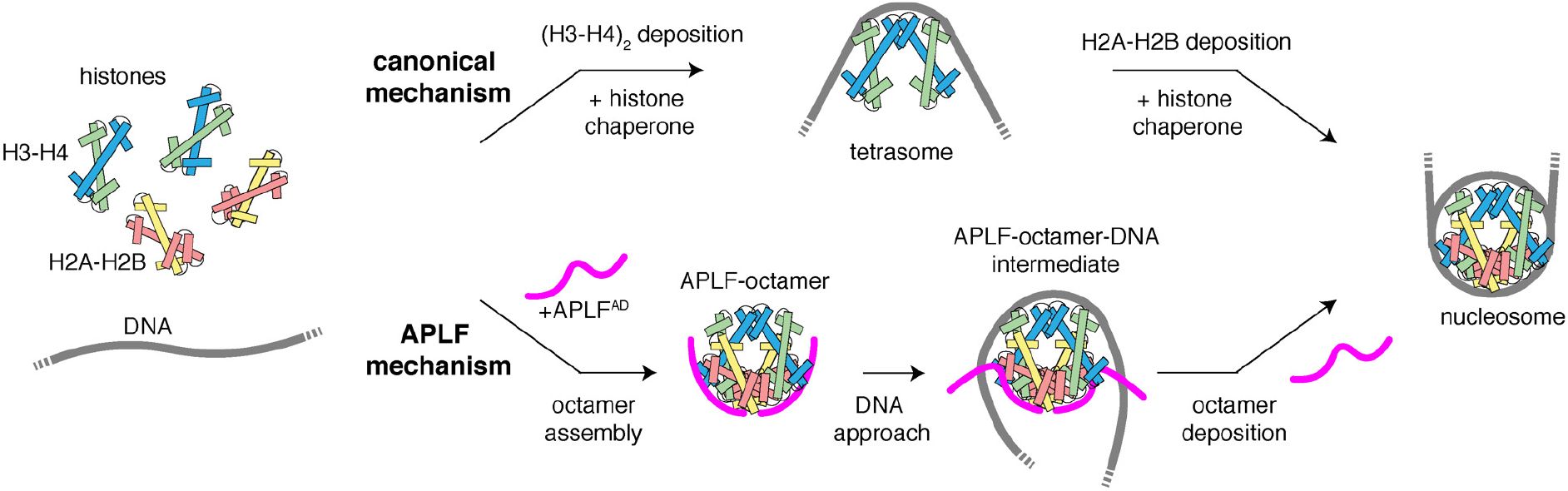
Model of the proposed histone chaperone mechanism by APLF. APLF^AD^ assembles H2A-H2B and H3-H4 simultaneously in an octameric configuration, and deposits them on DNA using a transient ternary intermediate. This contrasts with the stepwise nucleosome assembly where (H3-H4)^2^ deposition precedes H2A-H2B.

While many IDRs form fuzzy complexes where interactions are not well-defined, we find that APLF^AD^ forms specific and defined interactions that enable it to chaperone the histone octamer. Comparison of our data obtained for binding H2A-H2B alone (ref. (*14*)) and here on binding the histone octamer suggest that there can be a degree of fuzziness in the APLF^AD^-histone interaction, depending on the exact histone substrate: Y476 is important for high-affinity H2A-H2B binding but is not involved in histone octamer binding as judged from the crystal structure of the (APLF^AD-Δ^-H2A-H2B-H3-H4)_2_ complex. To what extent such adaptability is relevant for APLF function needs further investigation.

By tethering H2A-H2B and H3-H4 in their native nucleosomal configuration, APLF adds a remarkable new mode of action in the repertoire of histone chaperones. Our data demonstrates that histone octamer assembly can be uncoupled from nucleosome assembly and can be controlled by a single histone chaperone. Interestingly, while many histone chaperones contain acidic stretches(*32*), sequence analysis based on the presence of the key aromatic anchor residues revealed no clear candidates for a chaperone with similar histone octamer chaperone activity as APLF^AD^ (Fig. S16). It will be interesting to investigate whether some may nevertheless retain this function, and whether two histone chaperones may work together to chaperone the octamer by binding to each other, and how this is regulated. We speculate that chaperoning of octamers may be a more widespread mechanism to maintain H3-H4 as well as H2A-H2B with their modifications and variants within the same nucleosome, during temporary chromatin disassembly throughout the genome.

## Materials and Methods

### Data reporting

No statistical methods were used to predetermine sample size. The experiments were not randomized, and the investigators were not blinded to allocation during experiments and outcome assessment.

### Constructs, expression, and purification of APLF^AD^

The acidic domain of *Hs*. APLF (APLF^AD^, residues 450-511) was expressed and purified from a pLIC_His-GST-APLF^AD^ plasmid. Mutations were introduced using site-directed mutagenesis and verified by DNA sequencing. The quadruple mutant Y462A/F468A/Y476A/W485A (QA) and the truncated QA mutant (QA-Δ, residues 450-490), like all proteins for the chaperone assay, were expressed as fusion protein with a N-terminal SUMO-tag from a pET29b_SUMO-APLF^AD^ construct containing a His-tag and TEV cleavage site N-terminal to SUMO. Expression and purification were carried out as previously described with minor modifications(*14*). Briefly, APLF^AD^ (or SUMO-APLF^AD^) was expressed in BL21 Rosetta2 (DE3) cells (Novagen) at 30 °C. For NMR experiments, cells were cultured in M9 minimal medium in H_2_O containing ^15^NH_4_Cl for ^15^N-labeled APLF^AD^, or in D_2_O with ^15^NH_4_Cl and ^13^C_6_D_7_-glucose for perdeuterated ^15^N,^13^C-labeled APLF^AD^. After harvesting and lysis by freeze-thaw and sonification, soluble His-GST-APLF^AD^ (or His-SUMO-APLF^AD^) was loaded on a 5 mL HisTrap FF column (GE Healthcare Life Sciences), pre-equilibrated in lysis buffer (50 mM Tris, pH 8.0, 150 mM NaCl, 5 mM β - mercaptoethanol (BME), 20 mM imidazole), washed with lysis buffer, and eluted with a gradient of 20-500 mM imidazole in lysis buffer. The fusion protein was then cleaved with TEV protease (produced in-house) at 4 °C, typically overnight, and after complete cleavage, APLF^AD^ (or SUMO-APLF^AD^) was further purified by anion exchange on a 5 ml HiTrap Q HP column (GE Healthcare Life Sciences) in 20 mM Tris, pH 7.5, 5 mM BME, 1 mM EDTA with a salt gradient from 150 mM to 1 M NaCl. Fractions containing APLF^AD^ (or SUMO-APLF^AD^) were pooled, supplemented with MgCl_2_ (1.1 mM final concentration), and applied on a 5 mL HisTrap FF column (GE Healthcare Life Sciences) to remove residual His-tagged protein. The final purified APLF^AD^ (or SUMO-APLF^AD^) was pooled, buffer-exchanged to assay buffer (25 mM NaPi, pH 7.0, 300 mM NaCl) using a 3 kDa (or 10 kDa) molecular weight cut-off (MWCO) Amicon Ultra Centrifugal Filter Unit (Merck Millipore), and used directly or aliquoted, flash-frozen in liquid nitrogen and stored at -20 °C until further use.

### Histone production

Experiments were carried out with full-length *Drosophila melanogaster* (*Dm*) histones, except when noted otherwise. For crystallography, tailless *Xenopus laevis* (*Xl*) histones were used. Full-length *Dm*. histones H2A, H2B, H3, and H4 in pET21b plasmids and tailless *Xl*. H2A (residues 13-118), H2B (residues 24-122), and H4 (residues 20-102) in pET3a plasmids, and H3 (residues 38-135) in a pDEST plasmid were expressed in BL21 Rosetta2 (DE3) cells (Novagen) and purified from inclusion bodies as previously described except for minor modifications for the purification of tailless histones(*14, 33*). For NMR experiments on H3-H4, histone H3 was isotope-labeled by expression in M9 minimal medium in D_2_O with ^15^NH_4_Cl and ^13^C_6_D_7_-glucose. Briefly, after isolation of the inclusion bodies, solubilized histones were purified under denaturing conditions in two steps using size-exclusion and cation-exchange chromatography. First, histones were purified on a gel filtration column HiLoad Superdex 75 pg (GE Healthcare Life Sciences) pre-equilibrated with histone gel-filtration buffer (HGFB) (50 mM NaPi, pH 7.5, 5 mM BME, 1 mM EDTA, 7 M urea) with 150 mM NaCl (HGFB150) or, for tailless H4, 1 M NaCl (HGFB1000). Histone containing fractions were pooled and, for tailless histones H2A and H3, adjusted to a final NaCl concentration of 12.5 mM using HGFB, or, for tailless H4, using sodium acetate urea buffer (SAUB) (20 mM NaOAc, pH 5.2, 5 mM BME, 1 mM EDTA, 7 M urea). Histones were then further purified by cation exchange on a 5 ml HiTrap SP HP column (GE Healthcare Life Sciences), pre-equilibrated with HGFB150 (full-length histones and tailless H2B), HGFB (tailless H2A and H3) or SAUB (tailless H4). After a wash step, histones were eluted with a linear gradient of NaCl to 1 M in HGFB or SAUB. Histone containing fractions were pooled, concentrated, supplemented with 1 mM lysine (final concentration) and stored at -20 °C.

### 601-DNA production

A high-copy number plasmid containing 12 tandem repeats of a 167 base pair strong positioning DNA sequence (Widom’s 601(*34, 35*)) was transformed into DH5α cells. The plasmid was purified using a QIAGEN Plasmid Giga Kit. The 167-bp fragment was released from the vector by ScaI (Thermo Fisher Scientific) digestion and purified by anion exchange.

### Preparation of histone complexes

Histones were refolded and purified as previously described with minor changes(*14, 33*). Briefly, histone proteins were unfolded in 50 mM Tris, pH 7.5, 100 mM NaCl, 10 mM dithiothreitol (DTT), 6 M guanidine hydrochloride, mixed in equimolar ratios to a final protein concentration of 1 mg/ml, then dialyzed at 4 °C overnight to 10 mM Tris-HCl, pH 7.5, 1 mM EDTA, 5 mM BME, 2 M NaCl, followed by size-exclusion chromatography at 4 °C on a HiLoad Superdex 200 pg (GE Healthcare Life Sciences) column pre-equilibrated in the same buffer. For experiments using the stoichiometric core-histone mix (octamer-mix), the purified histone complexes were exchanged to assay buffer (25 mM NaPi, pH 7.0, 300 mM NaCl) using a 10 kDa MWCO Amicon Ultra Centrifugal Filter Unit (Merck Millipore). Complexes were aliquoted, flash frozen in liquid nitrogen and stored at -20 °C. Concentrations of H3-H4 are always given as concentration of dimers. Unless noted otherwise, concentrations of the octamer-mix are expressed as the equivalent histone octamer concentration, i.e., 10 µM octamer-mix equals 20 µM H2A-H2B and 20 µM H3-H4, corresponding to an equivalent histone octamer concentration of 10 µM.

### Analytical gel filtrations

Analytical gel filtrations were conducted in assay buffer (25 mM NaPi, pH 7.0, 300 mM NaCl) at room temperature. Histone complexes (20 µM H2A-H2B, 20 µM H3-H4, or 10 µM octamer-mix) were mixed with APLF^AD^, incubated for 30 min on ice, centrifuged to remove aggregates and then loaded on a Superdex 200 Increase 10/300 GL column (GE Healthcare Life Sciences) equilibrated in assay buffer and run at room temperature. Molar ratios of histone complex to APLF^AD^ ranged from 1:0 to 1:2 for H2A-H2B to APLF^AD^, 1:1.5 for H3-H4 to APLF^AD^, and 1:4 for histone octamer equivalent to APLF^AD^, as indicated in Fig. S3. The chromatogram in Fig. 1D is taken at 1:2 histone octamer equivalent to APLF^AD^.

### Isothermal titration calorimetry

Calorimetric titrations were conducted in assay buffer (25 mM NaPi, pH 7.0, 300 mM NaCl) at 25 °C using a MicroCal VP-ITC microcalorimeter (Malvern Panalytical). Calorimetric titrations of APLF^AD^ to H2A-H2B were described previously in ref. (*14*). For comparison between histone complexes, 10 µM H2A-H2B, 10 µM H3-H4 or 5 µM octamer-mix was used in the sample cell and titrated with 90 µM APLF^AD^ in the injection syringe. For binding comparison between H3-H4 and APLF^AD^ mutants, 20 µM H3-H4 in the cell was titrated with 180 µM APLF^AD^ in the syringe. For binding comparison between octamer-mix and APLF^AD^ mutants, 5 µM octamer-mix in the cell was titrated with 90 µM APLF^AD^ in the syringe. APLF^AD^ QA and QA-Δ mutants contained an N-terminal SUMO-tag for concentration determination. Comparison of ITC data on APLF^AD^ wild-type with and without SUMO-tag revealed little differences (Fig. S15). Binding isotherms were generated by plotting the heat change of the binding reaction against the ratio of total concentration of APLF^AD^ to total concentration of histone complexes. To allow direct comparison between H2A-H2B, H3-H4 and octamer-mix, the concentration of histone complexes was expressed as the total concentration of histone dimers, i.e., 5 µM octamer-mix corresponds to 10 µM H2A-H2B and 10 µM H3-H4, which equals 20 µM histone dimers. For comparison between APLF^AD^ mutants, the octamer-mix concentration was expressed as the equivalent histone octamer concentration. The enthalpy of binding (ΔH, kcal mol^-1^) was determined by integration of the injection peaks (5 µL) and correction for heats of dilution were determined from identical experiments without histone complexes. The entropy of binding (ΔS), the stoichiometry of binding (*n*), and the dissociation constant (*K*_D_) were determined by fitting the resulting corrected binding isotherms by nonlinear least-squares analysis to a one set of sites binding model using the Origin software (MicroCal, Inc.). Errors in fit parameters are the standard errors derived from the regression analyses as reported by the software.

### Native mass spectrometry

Complexes of histones and APLF^AD^ were prepared by mixing H2A-H2B or H3-H4 and APLF^AD^ in assay buffer (25 mM NaPi, pH 7.0, 300 mM NaCl) at a ratio of 1:1 histone dimer:APLF^AD^. The H2A-H2B-APLF^AD^ complex was purified on a Superdex 200 Increase 10/300 GL column (GE Healthcare Life Sciences) equilibrated in assay buffer and run at room temperature. Octamer-mix and APLF^AD^ were mixed in assay buffer at a ratio of equivalent to 1:0.25 to 1:3 histone octamer to APLF^AD^ and used without further purification. The mass spectrum in Fig. 1G is taken at 1:0.25 histone octamer equivalents to APLF^AD^. For each condition, a 20 µL sample at 20 µM concentration of complex (H2A-H2B– and H3-H4–APLF^AD^) or at 20 µM octamer-mix was buffer exchanged into 50 mM (H2A-H2B- and H3-H4-APLF^AD^) or 300 mM (octamer-mix + APLF^AD^) ammonium acetate at pH 7.5 using 3 kDa MWCO Amicon Ultra Centrifugal Filter Units (Merck Millipore). After buffer exchange the volume of each sample was ∼40 µL. The samples were then measured at the Exactive Plus EMR (Thermo Fisher Scientific) and the masses for each protein complex determined manually by minimization of the error over the charge state envelope from the different charge-state assignments.

### Cross-linking mass spectrometry

The stoichiometric core-histone mix (octamer-mix) and APLF^AD^ were mixed in assay buffer (25 mM NaPi, pH 7.0, 300 mM NaCl) at a ratio of equivalent to 1:0.25 to 1:2.5 histone octamer to APLF^AD^. The complex formed in the 1:2.5 mixture was purified on a Superdex 200 Increase 10/300 GL column (GE Healthcare Life Sciences) equilibrated in assay buffer and run at room temperature. The 1:0.25 mixture was used directly for mass-spectrometry. For each condition, 4 µL per reaction of 20 µM concentration of purified complex or 20 µM octamer-mix was diluted to 10 µM in 50 mM HEPES pH 7.5 and cross-linked for 15 minutes at room temperature with 500 µM disuccinimidyl sulfoxide (DSSO). The reaction was quenched with 1 M Tris pH 7.5 (50 mM final concentration). The cross-linking reaction was performed three times per sample. Each sample was supplemented with urea to 8 M, reduced by addition of DTT at a final concentration of 10 mM for 1 hour at room temperature, and alkylated for 0.5 hours at room temperature in the dark by addition of iodoacetamide at a final concentration of 50 mM, and quenched with DTT to 50 mM. The samples were digested in two rounds. In the first round, the samples were digested with Lys-C at an enzyme-to-protein ratio of 1:50 (w/w) at 30 °C for 3 hours, then diluted four times in 50 mM AmBic and further digested with trypsin at an enzyme-to-protein ratio of 1:100 (w/w) at 37 °C for 16 hours. The digested samples were desalted using homemade C18 stage tips, dried and stored at -80 °C until further use.

The samples were analyzed by LC-MS/MS using an Agilent 1290 Infinity System (Agilent Technologies) in combination with an Orbitrap Fusion Lumos (Thermo Scientific). Reverse phase chromatography was carried out using a 100-μm inner diameter 2-cm trap column (packed in-house with ReproSil-Pur C18-AQ, 3 μm) coupled to a 75-μm inner diameter 50 cm analytical column (packed in-house with Poroshell 120 EC-C18, 2.7 μm) (Agilent Technologies). Mobile-phase solvent A consisted of 0.1% formic acid in water, and mobile-phase solvent B consisted of 0.1% formic acid in 80% acetonitrile. A 120-minute gradient was used and start and end percentage of buffer B were adjusted to maximize sample separation. MS acquisition was performed using the MS2_MS3 strategy, where the MS1 scan was recorded in Orbitrap at a resolution of 60000, the selected precursors were fragmented in MS2 with CID and the cross-linker signature peaks recorded at a resolution of 30000, and the fragments displaying the mass difference specific for DSSO were further fragmented in a MS3 scan in the ion trap (IT)(*36*). All the samples were analyzed with Proteome Discoverer (version 2.2.0.388) with the XlinkX nodes integrated as described previously(*36, 37*).

For analysis, only cross-links reproduced in two out of three replicate experiments were considered. For analysis of intermolecular histone-histone cross-links, cross-links to the flexible tails of the histones were excluded as any cross-link with CMS < 1. The set of filtered inter-histone cross-links within the histone core were analyzed for compatibility with nucleosome structure (PDB 2PYO) by calculating the solvent accessible surface distance (SASD) between the Cα atoms of cross-linked lysines using Jwalk(*38*). Considering the maximum distance between the Cα atoms between DSSO cross-linked lysines is 23 Å, cross-links were categorized as incompatible with the native histone octamer structure when the SASD was more than 27 Å, using a 4 Å tolerance to account for backbone dynamics. Cross-links with SASD between 23 and 27 Å are only compatible when allowing for backbone dynamics while cross-links with SASD up to 23 Å are fully compatible with the native structure.

### NMR spectroscopy

All NMR experiments were carried out on Bruker Avance III HD spectrometers. NMR spectra were processed using Bruker TopSpin and analyzed using Sparky(*39*). NMR titrations of APLF^AD^ with H2A-H2B, H3-H4, or octamer-mix were done at 900 MHz ^1^H Larmor frequency at 298 K in NMR buffer (25 mM NaPi, pH 7, 5% D_2_O, with 1x protease inhibitors (complete EDTA-free cocktail, Roche)) with 600 mM NaCl for the titrations with H2A-H2B and H3-H4 and 300 mM NaCl for titration with octamer-mix. The titration was monitored using [^1^H-^15^N]-TROSY spectra, in 14 points from 1:0 to 1:2 molar ratio APLF^AD^:H2A-H2B and APLF^AD^:H3-H4 and in 4 points from 1:0 to 1:0.045 APLF^AD^ to histone octamer equivalent. For titrations with H2A-H2B and H3-H4, 20 µM of [U-^15^N]-APLF^AD^ was used, while for the titration with octamer-mix 300 µM [U-^2^H*/*^13^C/^15^N]-APLF^AD^ was used. Reported peak intensity ratios were corrected for differences in protein concentration (due to dilution) and number of scans. Residue-specific chemical shift perturbations (CSPs) were quantified from the perturbations in the ^1^H (Δδ_H_) and ^15^N (Δδ_N_) dimensions as the weighted average (composite) CSP in ppm:

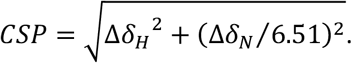

Assignment of H3 in H3-H4 was carried out using 325 µM [U-^2^H*/*^13^C/^15^N]-H3-H4 in 290 mM acetate buffer, pH 3.8, 1 mM EDTA, 5 mM BME, 5% D_2_O, 0.02% NaN_3_ with 1x protease inhibitors (complete EDTA-free cocktail, Roche). Backbone assignments were based on TROSY-based HNCA, HN(CO)CA, HNCB, and HN(CO)CB spectra, recorded at 900 MHz at 298 K. Backbone assignment was ∼80% complete and will be reported elsewhere. To map the binding site of APLF^AD^ on the H3-H4 surface, 20 µM [U-^2^H*/*^13^C/^15^N]H3-H4 was incubated in assay buffer (25 mM NaPi, pH 7.0, 300 mM NaCl) supplemented with 1 mM EDTA and 5 mM BME with 0, 10, or 20 µM peptide corresponding to APLF^459-474^ (sequence: Ac-PNEYDLNDSFLDDEEE-NH_2_, Biomatik, dissolved in assay buffer supplemented with 1 mM EDTA and 5 mM BME) for 30 minutes on ice. The mixture was subsequently buffer exchanged to 50 mM acetate buffer, pH 5, 1 mM EDTA, 5 mM BME, 5% D_2_O, 0.02% NaN_3_, with 1x protease inhibitors (complete EDTA-free cocktail, Roche) using a 10 kDa molecular weight cut-off (MWCO) Amicon Ultra Centrifugal Filter Unit (Merck Millipore). The titration was followed by [^1^H,^15^N]-TROSY spectra recorded at a 900 MHz spectrometer at 308 K.

To probe the interaction surface of APLF^AD^ for the octamer-mix, ^15^N TROSY CPMG relaxation dispersion experiments were performed at 298 K using [U-^2^H-^13^C-^15^N]APLF^AD^ alone or after the addition of 1.5% octamer-mix in NMR buffer with 300 mM NaCl. Data on the free protein were recorded at 900 MHz using a relaxation delay value of 40 ms (19 CPMG pulsing rates ranging between 25 and 1000 Hz including three replicates). CPMG dispersion profiles in presence of octamer-mix were recorded at 600 and 900 MHz using relaxation delay of, respectively, 40 and 20 ms (using 17 CPMG pulsing rates ranging between 25 and 1000 Hz for the 600 MHz data and 16 points between 50 and 1000 Hz, each time including three replicates). Peak intensities were extracted by fitting the line-shapes and converted to effective transverse relaxation rates, *R*_2,*eff*_, using PINT(*40, 41*). Dispersion profiles for resonances with significant dispersion of *R*_2,*eff*_ values (*R*_2,*eff*_ > 2 s^-1^ at 600 MHz) were subsequently fitted simultaneously in CATIA (https://www.ucl.ac.uk/hansen-lab/catia/) to extract chemical shift differences between free and bound states together with the population of the bound state and the exchange rate as global parameters using a two-site exchange model. Minimum error on *R*_2,*eff*_ during fitting was set to 2% or 0.3 s^-1^. Four residues (S467, L469, V497 and K511) were excluded from the final fit as their profiles indicated a more complex exchange behavior, resulting in a final data set of 28 dispersion profiles. The error surface of the fit was mapped using a grid-search, shown in Fig. S14.

### Small-angle X-ray scattering (SAXS)

Samples for SAXS were prepared in SAXS buffer (25 mM NaPi, pH 7, 300 mM NaCl, 3% v/v glycerol, 1 mM DTT). Octamer-mix was mixed with APLF^AD^ in assay buffer at a ratio corresponding to 1:2.5 histone octamer equivalents to APLF^AD^, concentrated using a 30 kDa MWCO Amicon Ultra Centrifugal Filter Unit (Merck Millipore), and purified on a HiLoad 16/600 Superdex 200 pg column (GE Healthcare Life Sciences) equilibrated in SAXS buffer and run at 4 °C. Elution fractions containing the complex were pooled, concentrated as above, flash-frozen in liquid nitrogen and stored at -20 °C until further use.

Synchrotron radiation X-ray scattering data from the complexes in size exclusion chromatography coupled SAXS (SEC-SAXS) and standard batch mode were collected at the EMBL P12 beamline of the storage ring PETRA III (DESY, Hamburg, Germany)(*42*). Images were collected using a photon counting Pilatus-6M detector at a sample to detector distance of 3.1 m and a wavelength (*λ*) of 0.12 nm covering the range of momentum transfer (*s*) 0.15 < s < 5 nm^-1^; with *s*=4πsin*θ*/*λ*, where 2*θ* is the scattering angle. In batch mode, a continuous flow cell capillary was used to reduce radiation damage. The latter was monitored by collecting 20 successive 50 ms exposures, comparing the frames, and discarding those displaying significant alterations. SEC-SAXS data was collected and analyzed as described previously(*43*). For this, a Superdex 200 incr. 10/300 column (GE Healthcare) was well equilibrated with SAXS buffer at a flow rate of 0.6 ml/min. SAXS data (3000 frames with 1 sec exposure) was collected on the sample after passing through the column. Data analysis was performed with Chromixs(*44*).

The final (background subtracted) SAXS profiles were subjected to standard SAXS analysis as follows. The data were normalized to the intensity of the transmitted beam and radially averaged; the scattering of pure buffer was used for background subtraction and the difference curves were scaled for solute concentration. The forward scattering *I*(0), the radius of gyration (*R*_g_) along with the probability distribution of the particle distances *P*(*r*) and the maximal dimension (*D*_max_) were computed using the automated SAXS data analysis pipeline SASFLOW(*45*) and various tools as implemented in ATSAS 2.8 package(*46*). The molecular masses (MM) were evaluated by comparison of the forward scattering with that from reference solutions of bovine serum albumin. In addition, several concentration-independent methods were applied utilizing empirical relationships between MM and several structural parameters obtained directly from the data(*47*). The *ab initio* bead modelling was performed using 10 independent runs of *DAMMIF*(*48*), from these the most probable model was selected for further analysis by *DAMAVER*(*49*). CRYSOL(*50*) was used to calculate the scattering profile from the atomic model described here and to compare with the experimental data.

### Chaperone assay

The chaperone assay was conducted in assay buffer (25 mM NaPi, pH 7.0, 300 mM NaCl) at room temperature. The ratio of octamer-mix to DNA (167 bp 601 sequence) that caused almost complete precipitation was determined experimentally at a ratio of 2–3 molar equivalents of histone octamer-mix to DNA. For the assay, octamer-mix (final reaction concentration: 2 µM) was pre-incubated alone or with APLF^AD^ wildtype (WT) (final reaction concentrations: 50, 100, 200 µM). All APLF^AD^ were with a N-terminally fused SUMO-tag. Binding of chaperone to histone was allowed to proceed at room temperature (RT) for 15 min before the addition of DNA to a final concentration of 1 µM in a total reaction volume of 20 µL. The reaction mixture was incubated at RT for 1 h followed by addition of 5 µL native PAGE loading buffer (10 mM Tris-HCl, pH 7.5, 1 mM EDTA, 1 mM DTT, 0.1 mM PMSF, 0.1 mg/ml BSA, 25% sucrose, 0.1% bromophenol blue) and removal of precipitates by centrifugation at 12,000 *g* for 5 minutes at 4 °C. Soluble complexes were separated on a pre-equilibrated 5% polyacrylamide gel, run in 0.2 × TBE (17.8 mM Tris, 17.8 mM boric acid, 0.4 mM EDTA) buffer at 150 V for 1 hour, at 4 °C. The gel was stained with DNA stain G (SERVA) before visualization using a Molecular Imager Gel Doc XR System (Bio-Rad).

### Nucleosome assembly assay and micrococcal nuclease digestion

Nucleosome assembly reactions were carried out as in the chaperone assay described above, with the following modifications: the reactions were run in 25 mM Tris pH 7, 300 mM NaCl (reaction buffer) using a 165 bp DNA fragment, all concentrated protein stocks were diluted in reaction buffer before use, concentration of octamer-mix was used at 3 µM, and the incubation was performed using untagged wild-type APLF^AD^ in a total reaction volume of 12 µL. After incubation of the octamer-mix with or without APLF^AD^ and DNA in reaction buffer, 4 µL of reaction mixture was transferred to fresh tubes and 1.25 µl 50% glycerol added to include as samples for native PAGE analysis before micrococcal nuclease (MNase) digestion. Another 5 µL from the reaction was transferred to fresh tubes to perform MNase digestion. The sample was diluted to a final volume of 25 µL and final buffer composition of 25 mM Tris pH 7, 150 mM NaCl. Each sample was mixed with 10 µL 10x MNase buffer (New England Biolabs), 1 µL 100x BSA (New England Biolabs), 1 µL of MNase (stock at 25 U/µL) (New England Biolabs) and 63 µl of water. After incubation at 37 °C for 10 minutes, the reactions were quenched by adding 10 µL 500 mM EDTA (final EDTA concentration ∼50 mM). The samples were treated with 25 µg Proteinase K (1.25 µL of 20 mg/mL stock solution, New England Biolabs) and incubated at 50 °C for 20 minutes. The MinElute PCR Purification Kit (Qiagen) was used to purify digested DNA fragments, after addition of a 621 bp loading control DNA fragment. The final elution was performed with 10 µL TE buffer (10 mM Tris pH 8, 1 mM EDTA). These samples, together with control samples taken before MNAse digestion, were run on a 6% polyacrylamide gel (Invitrogen) in 0.2 x TBE buffer at 150 V for 50 minutes, at RT. The gel was stained with SYBR™ Gold Nucleic Acid Gel Stain (Invitrogen) before visualization using a Molecular Imager Gel Doc XR System (Bio-Rad). Quantification of protected DNA fragments was performed using Bioanalyzer High sensitivity DNA chips, as previously described(*29*). The significance of the increase of bands in the nucleosomal size range was tested using one-tailed Students’ *t*-test in MATLAB 2016 (The MathWorks, Inc.).

### APLF-octamer-DNA ternary complex formation and detection

To test if the APLF^AD^-histone octamer complex can bind DNA, we used a native PAGE electrophoretic mobility shift assay using tetramethylrhodamine (TAMRA) fluorescent dye-tagged APLF, *Xenopus laevis* refolded histone octamer containing AlexaFluor647-labeled H2B T112C(*51*), and a 49 bp double-stranded DNA with the sequence GCACCGCTTAAACGCACGTACGCGCTGTCCCCCGCGTTTTAACCGCCAA (Eurofins) corresponding to the center of the 601 sequence(*34, 35*). Histones were obtained from histone source at Colorado State University (https://histonesource-colostate.nbsstore.net) and refolded as above. Labeled APLF^AD^ proteins were obtained by addition of TAMRA (16 mM) to APLF^AD^ (buffer exchanged to 50 mM NaPi, pH 8.3) at 2-4x molar excess of dye, followed by incubation over 2 days at 4 °C and purification using PD-10 Desalting Columns (Cytiva). Proteins were concentrated using 3 kDa MWCO Amicon Ultra-0.5 Centrifugal Filter Units (Merck Millipore). Protein concentrations were determined using the Proteins and Labels function on a Thermo Scientific™ NanoDrop™ One Microvolume UV-Vis Spectrophotometer and subsequently confirmed on SDS-PAGE gel upon Coomassie staining and colorimetric imaging on an Amersham ImageQuant 800 CCD Imager.

The APLF^AD^-histone octamer complex was assembled using the salt-dilution method from fluorescently tagged proteins. APLF^AD^ and histone octamer were mixed at 2:1 molar ratio in 25 mM NaPi, pH 7, 2 M NaCl and diluted stepwise by adding 25 mM NaPi, pH 7, no salt buffer to a final salt concentration of 0.6 M NaCl, targeting a final complex concentration of 20 µM. After removal of precipitates by spin-down at 12,000 g for 5 min and 4 °C, DNA was added at a 1:1 molar ratio to the histone octamer-chaperone complex (2 µM final concentration) and incubated for 1 hour at room temperature in 25 mM NaPi, pH 7, 300 mM NaCl. The ternary complex was obtained after centrifugation at 12,000 g for 5 min and 4 °C to remove precipitates. The ternary complex was crosslinked using disuccinimidyl suberate (DSS) (Thermo Fisher Scientific) at 1 mM for 20 min at room temperature. Control reactions used DMSO instead of the crosslinker. Samples were centrifuged at 12,000 g for 5 min and 4 °C before loading with 20% glycerol on a 6% DNA retardation polyacrylamide gel, run in 0.2x TBE at 120 V for 1 hour at 4 °C. Gels were imaged on an Amersham ImageQuant 800 CCD Imager, using a Cy3 scan for APLF signal and a Cy5 scan for the histone octamer. After SYBR Gold staining, the Cy3 scan detected predominantly the DNA signal with a small contribution of the APLF signal and the Cy5 scan detected histone octamer signal. Scans were merged using ImageJ software(*52*).

### Crystallization and data collection

For crystallization attempts, various APLF and histone constructs were tested. Initial conditions with full-length proteins led to clear drops and phase separation. Crystals used for structure determination were obtained with APLF^AD-Δ^ (*Homo sapiens* (*Hs*.) APLF residues 449-490, with N-terminal Gly as leftover from TEV-cleavage site) and tailless *Xenopus laevis* (*Xl*.) histones H2A (residues 13-118), H2B (residues 27-125), H3 (residues 38-135), and H4 (residues 20-102). To reconstitute the complex, APLF^AD-Δ^ was buffer exchanged to 10 mM Tris-HCl, pH 7.5, 1 mM EDTA, 5 mM BME, 2 M NaCl using a 3 kDa MWCO Amicon Ultra Centrifugal Filter Unit (Merck Millipore) and mixed with tailless *Xl* histone octamer in the same buffer at a molar ratio 2:1 APLF^AD-Δ^:histone octamer on ice. After stepwise dilution to 600 mM NaCl, the complex was concentrated using a 30 kDa MWCO Amicon Ultra Centrifugal Filter Unit (Merck Millipore) and purified by gel filtration on a HiLoad 16/600 Superdex 200 pg column (GE Healthcare Life Sciences) pre-equilibrated with 20 mM HEPES, pH 7.5, 1 mM DTT, 600 mM NaCl. Elution fractions containing the complex were pooled and concentrated as above and used directly for crystal screening using the vapor diffusion sitting drop method. Crystals used for the structure determination were obtained from the commercial screen JCSG+ Suite (Qiagen) by mixing complex and reservoir solution in an MRC2 plate (SWISSCI) at two different ratios (1.25:0.75 and 0.75:1.25 complex:reservoir). Crystals grew in up to 4 months and at 20 °C in a solution of 0.1 M sodium cacodylate pH 6.5, 1 M tri-sodium citrate. Crystals were transferred to the reservoir solution supplemented with 20% glycerol and quickly frozen in liquid nitrogen. Crystallographic data were collected on beamline X6A at the Swiss Light Source and the structure was refined to 2.35 Å resolution (Extended Data Table 1).

### Crystal structure determination, model building and refinement

All data were processed and scaled with XDS package and Aimless(*53, 54*). Structure determination was performed by molecular replacement with Phaser(*55*), using the H2A-H2B-H3-H4 octamer structure (PDB: 2HIO) as searching model. Model building and refinement were performed with COOT(*56*), phenix refine(*57*) and PDB-REDO(*58*). The data collection and refinement statistics are summarized in Extended Data Table 1. There are four NCS complexes in the asymmetric unit. Depending slightly on the copy, the final model includes residues 16/17 to 118/119 for H2A, 36/37 to 124/125 for H2B, 41 to 135 for H3, 25/27 to 100/101 for H4, and 458/459 to 487 for APLF^AD^, with clear density for APLF^AD^ residues 472/474 to 479/481 missing in all but one APLF^AD^ chain (see Fig. S7). The continuous density for one APLF^AD^ chain is likely the result of a crystal packing interaction as shown in Fig. S10. Figures were made using PYMOL (the PyMOL Molecular Graphics System, version 2.3, Schrödinger, LLC). Plots of electrostatic surface potential were generated using the APBS-tool(*59*) in PyMOL.

### Structural modelling of the full-length APLF^AD^ complex

The model of the full-length APLF^AD^ bound to histone octamer was derived from the APLF^AD-Δ^-histone octamer crystal structure by first building the missing APLF^AD^ residues using MODELLER(*60*), and then refining the resulting models in HADDOCK(*61*). Briefly, first the missing residues in the linker region (475-481) of one APLF^AD^ chain were added, selecting the best ranking model for further modelling. Second, the missing N-terminal residues (450-458) were built, selecting the model with least contact between the N-terminal segment and the histone surface, in line with the NMR results that show that the N-terminal residues remain highly flexible in the complex. Third, the two missing C-terminal H2B residues were built, selecting the best ranking model. Fourth, the missing C-terminal APLF^AD^ residues were built taking into account the intermolecular cross-links between APLF^AD^ and the histones observed at 1:1:0.125 H2A-H2B:H3-H4:APLF^AD^. For each APLF^AD^ chain, two distance restraints were added with 25 Å upper limit and 1 Å tolerance to restrain the Cα-Cα distance of the cross-linked residues (APLF^AD^ K505 to H2B K122 and APLF^AD^ K509 to H2B K113). Since the experimental data do not discriminate which copy of APLF^AD^ is cross-linked to which copy of H2B, two modelling runs were performed. In the first, the cross-links were set between the APLF^AD^ and the H2B copy that is already bound by the same APLF^AD^ copy (the proximal H2B). In the second run, the cross-links were set between the APLF^AD^ and the H2B copy that is bound by the other APLF^AD^ copy (the distal H2B). In each run 20 models were built. From each model, the APLF^AD^ coordinates were extracted, resulting in total in 40 conformations for each APLF^AD^ chain. Next, these models were refined in HADDOCK to select the final ensemble of 20 best solutions. Docking was set up as a three-body docking with the histone octamer structure and one ensemble of 20 conformations (either of the H2B distal or H2B proximal variety) for each APLF^AD^ chain. The starting structures were fixed to original position in the rigid body docking phase. Of the 400 models generated, 200 were refined in the semi-flexible refinement stage and subsequently refined in explicit water. From the 50 best ranking structures from the H2B distal and the H2B proximal run, those structures in which the solvent accessible surface distance (SASD) between the Cα atoms of cross-linked lysines, calculated using Jwalk(*38*), was less than 27 Å were extracted, combined and sorted using their HADDOCK score. The 20 best scoring models were selected as the final ensemble. This ensemble contained 7 APLF^AD^ in the H2B proximal conformation and 13 in the H2B distal conformation.

### Cell line, transfections, and plasmids

U2OS cells were cultured in 5% CO_2_ at 37 °C in DMEM (Dulbecco’s modified Eagle’s medium) supplemented with 10% fetal calf serum and antibiotics. U2OS cells were transfected with plasmid DNA using Lipofectamine 2000 (Invitrogen) according to the manufacturer’s instructions and analysed 24 hours after transfection. The expression vector for full-length human APLF was amplified from plasmid APLF-PC1-PURO’(*62*) and cloned into pCDNA5/FRT/TO-Puro as a XhoI*/HindIII* fragment using primers listed in Supplementary Table S4. APLF mutants DA-34 (Y462A/F468A), DA-AB (Y476A/W485A), QA (Y462A/F468A/Y476A/W485A) and ΔAD were generated by site-directed mutagenesis PCR using primers listed in Supplementary Table S4. All APLF expression constructs were verified using Sanger sequencing. The plasmid for mCherry-NBS1 expression was previously described(*63*). Generation of U2OS Flp-In/T-Rex cells for eGFP-APLF expression was previously described(*64*). Briefly, pCDNA5/FRT/TO-Puro plasmid encoding eGFP-APLF wildtype and mutants (5 µg), were co-transfected together with pOG44 plasmid encoding the Flp recombinase (1 µg). After selection on 1 µg/mL puromycin, single clones were isolated and expanded. U2OS Flp-In/T-Rex clones were incubated with 2 µg/mL doxycycline for 24 h to induce expression of cDNAs.

### GFP Pull-down assays

GFP pull-downs were performed on U2OS Flp-In/T-Rex cells expressing eGFP-APLF-WT or the indicated eGFP-tagged APLF mutants as previously described(*64*). Cells were lysed in EBC buffer (50 mM Tris, pH 7.5, 150 mM NaCl, 0.5% NP-40, 1 mM MgCl_2_, protease inhibitor cocktail tablets) with 500 units benzonase. Samples were incubated for 1 h at 4 °C under constant mixing. 50 μL input sample was collected in a separate tube and mixed with 2× Laemmli buffer. The cleared lysates were subjected to GFP pull-down with GFP-Trap beads (Chromotek). The beads were then washed six times with EBC buffer and boiled in 2× Laemmli buffer along with the input samples. Samples were subjected to western blot analysis using primary antibodies listed in Table S5.

### 365 nm UV-A Laser micro-irradiation and APLF recruitment

U2OS cells were grown on 18-mm coverslips and sensitized with 15 µM 5’-bromo-2-deoxyuridine (BrdU) for 24 h before micro-irradiation. For micro-irradiation, cells were placed in a live-cell imaging chamber set to 37 °C in CO_2_-independent Leibovitz’s L15 medium supplemented with 10% FCS. Live cell imaging and micro-irradiation experiments were carried out with a Zeiss Axio Observer microscope driven by ZEN software using a ×63/1.4 oil immersion objective coupled to a 355 nm pulsed DPSS UV-laser (Rapp OptoElectronic). Images were recorded using ZEN 2012 software and analyzed in Image J(*52*). The integrated density of laser tracks was measured within the locally irradiated area (*I*_damage_) and divided over that area. The same was done for the nucleoplasm outside the locally irradiated area (*I*_nucleoplasm_) and in a region not containing cells in the same field of view (*I*_background_). The level of protein accumulation in the laser track relative to the protein level in the nucleoplasm was calculated as follows: ((*I*_damage_ − *I*_background_)/(*I*_nucleoplasm_ − *I*_background_) − 1).

## Acknowledgements

We thank Raymond Schellevis for assistance in protein purification, Gert Folkers for lab support, and Johan van der Zwan and Andrei Gurinov for support and maintenance of the NMR infrastructure (all from the Utrecht NMR Group at Utrecht University). We thank Patrick Celie and Alexander Fish for help in setting up crystallization screens and Tatjana Heidebrecht for fishing crystals (all from Netherlands Cancer Institute). We thank the Swiss Light Source staff for X-ray diffraction synchrotron access and support. We thank the beamline P12 staff of EMBL Hamburg and the PETRA III storage ring staff of DESY (Hamburg, Germany) for SAXS synchrotron access. We thank Anneloes Blok and Marcellus Ubbink from Macromolecular Biochemistry, Leiden Institute of Chemistry, Leiden University for providing access to the VP-ITC MicroCalorimeter. We thank Pamela Dyer and Karolin Luger from the Department of Biochemistry, University of Colorado at Boulder for providing plasmids of tailless histones H3 and H4.

## Funding

Marie Curie Initial Training Network Innovative Doctoral Programme ManiFold grant 317371 EC-FP7/2007-201 to the Utrecht University Bijvoet Centre.

Netherlands Organization for Scientific Research grant 723.013.010 (HvI)

European Research Council grant 851564 (FM)

Netherlands Organization for Scientific Research grant 175.107.301.10 (RB)

Netherlands Organization for Scientific Research grant 700.53.103 (RB)

Netherlands Organization for Scientific Research National Roadmap grant 184.032.207 to uNMR-NL, the National Roadmap Large-Scale NMR Facility of the Netherlands

European Research Council grant 731005 to INSTRUCT, for Instruct-ULTRA, an EU H2020 project to further develop the services of Instruct-ERIC

European Research Council project 823839 to Epic-X, an EU Horizon 2020 program INFRAIA project.

European Research Council grant 653706 to INEXT and grant 871037 to iNEXT-Discovery, an EU Horizon 2020 program, which provided financial support for SAXS measurements at beamline P12 operated by EMBL Hamburg at the PETRA III storage ring (DESY, Hamburg, Germany)

## Author contributions

Conceptualization: IC, FM, TKS, HVI

Investigation: IC, XG, BVE, FM, DF, MAG, WW, KV

Visualization: IC, FM, HVI

Funding acquisition: HVI, RB

Project administration: HVI, RB

Supervision: HVI, RB, HW, FM, TS, HVA, AH, TKS, DS

Writing – original draft: IC, HVI with input from all authors

Writing – review & editing: IC, FM, HVI with input from all authors

## Data availability

All data needed to evaluate the conclusions in the paper are present in the paper and/or the Supplementary Materials. The coordinates of the APLF^AD^-histone octamer complex have been deposited in the Protein Data Bank under accession number 6YN1. The experimental SAXS data and models are deposited in SASBDB with the accession codes: SASDJJ5. The Native MS and cross-linking mass spectrometry data are available via Figshare: https://figshare.com/s/43e763c4f078ce450cf8 (temporary reviewer link).

## Supplementary Materials

Figs. S1 to S16

Tables S1 to S5

## References

1. R. A. Laskey, B. M. Honda, A. D. Mills, J. T. Finch, Nucleosomes are assembled by an acidic protein which binds histones and transfers them to DNA. Nature. 275, 416–420 (1978).

2. C. M. Hammond, C. B. Strømme, H. Huang, D. J. Patel, A. Groth, Histone chaperone networks shaping chromatin function. Nat. Rev. Mol. Cell Biol. 18, 141–158 (2017).

3. Z. A. Gurard-Levin, J.-P. Quivy, G. Almouzni, Histone Chaperones: Assisting Histone Traffic and Nucleosome Dynamics. Annu. Rev. Biochem. 83, 487–517 (2014).

4. J. A. Kleinschmidt, A. Seiter, H. Zentgraf, Nucleosome assembly in vitro: Separate histone transfer and synergistic interaction of native histone complexes purified from nuclei of Xenopus laevis oocytes. EMBO J. 9, 1309–1318 (1990).

5. S. J. Elsässer, S. D’Arcy, Towards a mechanism for histone chaperones. Biochim. Biophys. Acta -Gene Regul. Mech. 1819, 211–221 (2012).

6. N. Iles, S. Rulten, S. F. El-Khamisy, K. W. Caldecott, APLF (C2orf13) Is a Novel Human Protein Involved in the Cellular Response to Chromosomal DNA Strand Breaks. Mol. Cell. Biol. 27, 3793–3803 (2007).

7. C. J. Macrae, R. D. McCulloch, J. Ylanko, D. Durocher, C. A. Koch, APLF (C2orf13) facilitates nonhomologous end-joining and undergoes ATM-dependent hyperphosphorylation following ionizing radiation. DNA Repair (Amst). 7, 292–302 (2008).

8. G. J. Grundy, S. L. Rulten, Z. Zeng, R. Arribas-Bosacoma, N. Iles, K. Manley, A. Oliver, K. W. Caldecott, APLF promotes the assembly and activity of non-homologous end joining protein complexes. EMBO J. 32, 112–125 (2013).

9. L. Woodbine, A. R. Gennery, P. A. Jeggo, The clinical impact of deficiency in DNA non-homologous end-joining. DNA Repair (Amst). 16, 84–96 (2014).

10. B. J. Sishc, A. J. Davis, The role of the core non-homologous end joining factors in carcinogenesis and cancer. Cancers (Basel). 9 (2017).

11. M. Hammel, Y. Yu, S. K. Radhakrishnan, C. Chokshi, M. S. Tsai, Y. Matsumoto, M. Kuzdovich, S. G. Remesh, S. Fang, A. E. Tomkinson, S. P. Lees-Miller, J. A. Tainer, An intrinsically disordered APLF links Ku, DNA-PKcs, and XRCC4-DNA ligase IV in an extended flexible non-homologous end joining complex. J. Biol. Chem. 291, 26987–27006 (2016).

12. P. Shirodkar, A. L. Fenton, L. Meng, C. A. Koch, Identification and functional characterization of a Ku-binding motif in aprataxin polynucleotide kinase/phosphatase-like factor (APLF). J. Biol. Chem. 288, 19604–19613 (2013).

13. P. V. Mehrotra, D. Ahel, D. P. Ryan, R. Weston, N. Wiechens, R. Kraehenbuehl, T. Owen-Hughes, I. Ahel, DNA Repair Factor APLF Is a Histone Chaperone. Mol. Cell. 41, 46–55 (2011).

14. I. Corbeski, K. Dolinar, H. Wienk, R. Boelens, H. Van Ingen, DNA repair factor APLF acts as a H2A-H2B histone chaperone through binding its DNA interaction surface. Nucleic Acids Res. 46, 7138–7152 (2018).

15. S. Dutta, I. V. Akey, C. Dingwall, K. L. Hartman, T. Laue, R. T. Nolte, J. F. Head, C. W. Akey, The crystal structure of nucleoplasmin-core: Implications for histone binding and nucleosome assembly. Mol. Cell. 8, 841–853 (2001).

16. K. F. Tóth, J. Mazurkiewicz, K. Rippe, Association states of nucleosome assembly protein 1 and its complexes with histones. J. Biol. Chem. 280, 15690–15699 (2005).

17. S. Muto, M. Senda, Y. Akai, L. Sato, T. Suzuki, R. Nagai, T. Senda, M. Horikoshi, Relationship between the structure of SET/TAF-Iβ/INHAT and its histone chaperone activity. Proc. Natl. Acad. Sci. U. S. A. 104, 4285–4290 (2007).

18. Y. Tsunaka, Y. Fujiwara, T. Oyama, S. Hirose, K. Morikawa, Integrated molecular mechanism directing nucleosome reorganization by human FACT. Genes Dev. 30, 673–686 (2016).

19. M. Zhang, H. Liu, Y. Gao, Z. Zhu, Z. Chen, P. Zheng, L. Xue, J. Li, M. Teng, L. Niu, Structural Insights into the Association of Hif1 with Histones H2A-H2B Dimer and H3-H4 Tetramer. Structure. 24, 1810–1820 (2016).

20. Y. Lorch, M. Zhang, R. D. Kornberg, Histone octamer transfer by a Chromatin-Remodeling Complex. 96, 389–392 (1999).

21. C. E. Rowe, G. J. Narlikar, The ATP-dependent remodeler RSC transfers histone dimers and octamers through the rapid formation of an unstable encounter intermediate. Biochemistry. 49, 9882–9890 (2010).

22. J. Markert, K. Zhou, K. Luger, SMARCAD1 is an ATP-dependent histone octamer exchange factor with de novo nucleosome assembly activity. Sci. Adv. 7, 1–12 (2021).

23. K. Luger, A. W. Mäder, R. K. Richmond, D. F. Sargent, T. J. Richmond, Crystal structure of the nucleosome core particle at 2.8 Å resolution. Nature. 389, 251–260 (1997).

24. C. A. Davey, D. F. Sargent, K. Luger, A. W. Maeder, T. J. Richmond, Solvent Mediated Interactions in the Structure of the Nucleosome Core Particle at 1.9 Å Resolution. J. Mol. Biol. 319, 1097–1113 (2002).

25. D. J. Kemble, L. L. McCullough, F. G. Whitby, T. Formosa, C. P. Hill, FACT Disrupts Nucleosome Structure by Binding H2A-H2B with Conserved Peptide Motifs. Mol. Cell. 60, 294–306 (2015).

26. Y. Wang, S. Liu, L. Sun, N. Xu, S. Shan, F. Wu, X. Liang, Y. Huang, E. Luk, C. Wu, Z. Zhou, Structural insights into histone chaperone Chz1-mediated H2A.Z recognition and histone replacement. PLoS Biol. 17, 1–20 (2019).

27. M. D. Ricketts, J. Han, M. R. Szurgot, R. Marmorstein, Molecular basis for chromatin assembly and modification by multiprotein complexes. Protein Sci. 28, 329–343 (2019).

28. M. Hondele, T. Stuwe, M. Hassler, F. Halbach, A. Bowman, E. T. Zhang, B. Nijmeijer, C. Kotthoff, V. Rybin, S. Amlacher, E. Hurt, A. G. Ladurner, Structural basis of histone H2A-H2B recognition by the essential chaperone FACT. Nature. 499, 111–114 (2013).

29. F. Mattiroli, Y. Gu, K. Luger, Measuring Nucleosome Assembly Activity in vitro with the Nucleosome Assembly and Quantification (NAQ) Assay. Bio-Protocol. 8, 1–11 (2018).

30. S. Smith, B. Stillman, Stepwise assembly of chromatin during DNA replication in vitro. EMBO J. 10, 971–980 (1991).

31. M. A. Hall, A. Shundrovsky, L. Bai, R. M. Fulbright, J. T. Lis, M. D. Wang, High-resolution dynamic mapping of histone-DNA interactions in a nucleosome. Nat. Struct. Mol. Biol. 16, 124–129 (2009).

32. C. Warren, D. Shechter, Fly Fishing for Histones : Catch and Release by Histone Chaperone Intrinsically Disordered Regions and Acidic Stretches. J. Mol. Biol. 429, 2401–2426 (2017).

33. K. Luger, T. J. Rechsteiner, T. J. Richmond, Preparation of nucleosome core particle from recombinant histones. Methods Enzymol. 304, 3–19 (1999).

34. P. T. Lowary, J. Widom, New DNA sequence rules for high affinity binding to histone octamer and sequence-directed nucleosome positioning. J. Mol. Biol. 276, 19–42 (1998).

35. A. ThÅström, L. M. Bingham, J. Widom, Nucleosomal Locations of Dominant DNA Sequence Motifs for Histone–DNA Interactions and Nucleosome Positioning. J. Mol. Biol. 338, 695–709 (2004).

36. F. Liu, P. Lössl, R. Scheltema, R. Viner, A. J. R. Heck, Optimized fragmentation schemes and data analysis strategies for proteome-wide cross-link identification. Nat. Commun. 8 (2017).

37. O. Klykov, B. Steigenberger, S. Pektaş, D. Fasci, A. J. R. Heck, R. A. Scheltema, Efficient and robust proteome-wide approaches for cross-linking mass spectrometry. Nat. Protoc. 13, 2964–2990 (2018).

38. J. M. A. Bullock, J. Schwab, K. Thalassinos, M. Topf, The importance of non-accessible crosslinks and solvent accessible surface distance in modeling proteins with restraints from crosslinking mass spectrometry. Mol. Cell. Proteomics. 15, 2491–2500 (2016).

39. W. Lee, M. Tonelli, J. L. Markley, NMRFAM-SPARKY: Enhanced software for biomolecular NMR spectroscopy. Bioinformatics. 31, 1325–1327 (2015).

40. M. Niklasson, R. Otten, A. Ahlner, C. Andresen, J. Schlagnitweit, K. Petzold, P. Lundström, Comprehensive analysis of NMR data using advanced line shape fitting. J. Biomol. NMR. 69, 93–99 (2017).

41. A. Ahlner, M. Carlsson, B. H. Jonsson, P. Lundström, PINT: A software for integration of peak volumes and extraction of relaxation rates. J. Biomol. NMR. 56, 191–202 (2013).

42. C. E. Blanchet, A. Spilotros, F. Schwemmer, M. A. Graewert, A. Kikhney, C. M. Jeffries, D. Franke, D. Mark, R. Zengerle, F. Cipriani, S. Fiedler, M. Roessle, D. I. Svergun, Versatile sample environments and automation for biological solution X-ray scattering experiments at the P12 beamline (PETRA III, DESY). J. Appl. Crystallogr. 48, 431–443 (2015).

43. M. A. Graewert, S. Da Vela, T. W. Gräwert, D. S. Molodenskiy, C. E. Blanchet, D. I. Svergun, C. M. Jeffries, Adding Size Exclusion Chromatography (SEC) and Light Scattering (LS) Devices to Obtain High-Quality Small Angle X-Ray Scattering (SAXS) Data. Cryst.. 10 (2020).

44. A. Panjkovich, D. I. Svergun, CHROMIXS: Automatic and interactive analysis of chromatography-coupled small-angle X-ray scattering data. Bioinformatics. 34, 1944–1946 (2018).

45. D. Franke, A. G. Kikhney, D. I. Svergun, Automated acquisition and analysis of small angle X-ray scattering data. Nucl. Instruments Methods Phys. Res. Sect. A Accel. Spectrometers, Detect. Assoc. Equip. 689, 52–59 (2012).

46. D. Franke, M. V. Petoukhov, P. V. Konarev, A. Panjkovich, A. Tuukkanen, H. D. T. Mertens, A. G. Kikhney, N. R. Hajizadeh, J. M. Franklin, C. M. Jeffries, D. I. Svergun, ATSAS 2.8: A comprehensive data analysis suite for small-angle scattering from macromolecular solutions. J. Appl. Crystallogr. 50, 1212–1225 (2017).

47. N. R. Hajizadeh, D. Franke, C. M. Jeffries, D. I. Svergun, Consensus Bayesian assessment of protein molecular mass from solution X-ray scattering data. Sci. Rep. 8, 1–13 (2018).

48. D. Franke, D. I. Svergun, DAMMIF, a program for rapid ab-initio shape determination in small-angle scattering. J. Appl. Crystallogr. 42, 342–346 (2009).

49. V. V. Volkov, D. I. Svergun, Uniqueness of ab initio shape determination in small-angle scattering. J. Appl. Crystallogr. 36, 860–864 (2003).

50. D. Svergun, C. Barberato, M. H. Koch, CRYSOL - A program to evaluate X-ray solution scattering of biological macromolecules from atomic coordinates. J. Appl. Crystallogr. 28, 768–773 (1995).

51. U. Muthurajan, F. Mattiroli, S. Bergeron, K. Zhou, Y. Gu, S. Chakravarthy, P. Dyer, T. Irving, K. Luger, In Vitro Chromatin Assembly: Strategies and Quality Control. Methods Enzymol. 573, 3–41 (2016).

52. C. A. Schneider, W. S. Rasband, K. W. Eliceiri, NIH Image to ImageJ: 25 years of image analysis. Nat. Methods. 9, 671–675 (2012).

53. W. Kabsch, Xds. Acta Crystallogr. Sect. D Biol. Crystallogr. 66, 125–132 (2010).

54. P. R. Evans, G. N. Murshudov, How good are my data and what is the resolution? Acta Crystallogr. Sect. D Biol. Crystallogr. 69, 1204–1214 (2013).

55. A. J. McCoy, R. W. Grosse-Kunstleve, P. D. Adams, M. D. Winn, L. C. Storoni, R. J. Read, Phaser crystallographic software. J. Appl. Crystallogr. 40, 658–674 (2007).

56. P. Emsley, B. Lohkamp, W. G. Scott, K. Cowtan, Features and development of Coot. Acta Crystallogr. Sect. D Biol. Crystallogr. 66, 486–501 (2010).

57. P. V. Afonine, R. W. Grosse-Kunstleve, N. Echols, J. J. Headd, N. W. Moriarty, M. Mustyakimov, T. C. Terwilliger, A. Urzhumtsev, P. H. Zwart, P. D. Adams, Towards automated crystallographic structure refinement with phenix. refine. Acta Crystallogr. Sect. D Biol. Crystallogr. 68, 352–367 (2012).

58. R. P. Joosten, J. Salzemann, V. Bloch, H. Stockinger, A. C. Berglund, C. Blanchet, E. Bongcam-Rudloff, C. Combet, A. L. Da Costa, G. Deleage, M. Diarena, R. Fabbretti, G. Fettahi, V. Flegel, Gisel, V. Kasam, T. Kervinen, E. Korpelainen, K. Mattila, M. Pagni, M. Reichstadt, V. Breton, J. Tickle, G. Vriend, PDB-REDO: Automated re-refinement of X-ray structure models in the PDB. J. Appl. Crystallogr. 42, 376–384 (2009).

59. N. A. Baker, D. Sept, S. Joseph, M. J. Holst, J. A. McCammon, Electrostatics of nanosystems: Application to microtubules and the ribosome. Proc. Natl. Acad. Sci. U. S. A. 98, 10037–10041 (2001).

60. B. Webb, A. Sali, Comparative protein structure modeling using MODELLER. Curr. Protoc. Protein Sci. 86, 2.9.1–2.9.37 (2016).

61. G. C. P. Van Zundert, J. P. G. L. M. Rodrigues, M. Trellet, C. Schmitz, P. L. Kastritis, E. Karaca, A. S. J. Melquiond, M. Van Dijk, S. J. De Vries, A. M. J. J. Bonvin, The HADDOCK2.2 Web Server: User-Friendly Integrative Modeling of Biomolecular Complexes. J. Mol. Biol. 428, 720–725 (2016).

62. S. L. Rulten, F. Cortes-Ledesma, L. Guo, N. J. Iles, K. W. Caldecott, APLF (C2orf13) is a novel component of poly(ADP-ribose) signaling in mammalian cells. Mol. Cell. Biol. 28, 2620–2628 (2008).

63. M. S. Luijsterburg, I. de Krijger, W. W. Wiegant, R. G. Shah, G. Smeenk, A. J. L. de Groot, A. Pines, A. C. O. Vertegaal, J. J. L. Jacobs, G. M. Shah, H. van Attikum, PARP1 Links CHD2-Mediated Chromatin Expansion and H3.3 Deposition to DNA Repair by Non-homologous End-Joining. Mol. Cell. 61, 547–562 (2016).

64. J. K. Singh, R. Smith, M. B. Rother, A. J. L. de Groot, W. W. Wiegant, K. Vreeken, O. D’Augustin, R. Q. Kim, H. Qian, P. M. Krawczyk, R. González-Prieto, A. C. O. Vertegaal, M. Lamers, S. Huet, H. van Attikum, Zinc finger protein ZNF384 is an adaptor of Ku to DNA during classical non-homologous end-joining. Nat. Commun. 12, 1–21 (2021).

