## Supplementary Materials for "Chaperoning of the histone octamer by the acidic domain of DNA repair factor APLF"

### Supplementary Figures

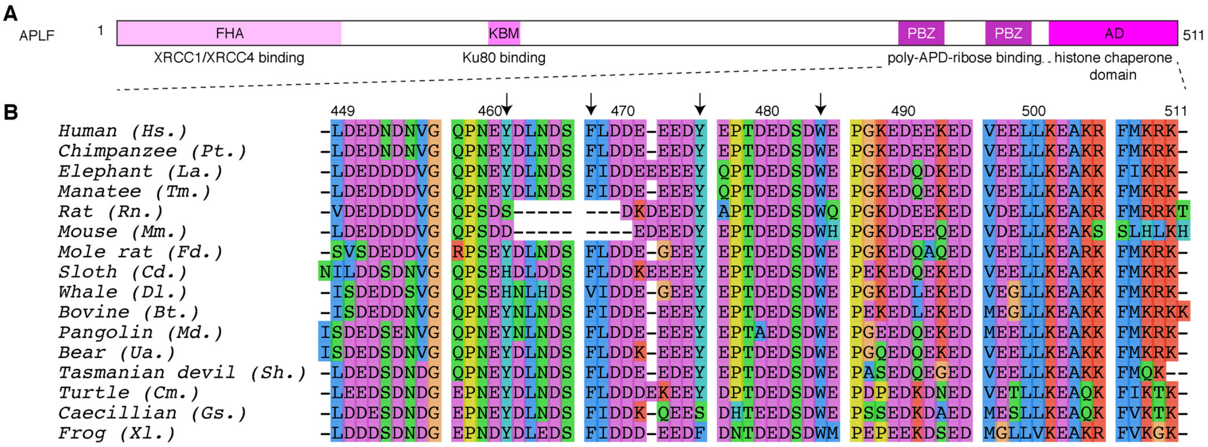


**Fig. S1 | Sequence conservation of the APLF acidic domain. A**, Domain organization of APLF, indicating the position of the forkhead associated (FHA) domain that binds XRCC4, the Ku-binding motif (KBM) that binds Ku80, the poly-ADP-ribose binding zinc-finger (PBZ) domains that bind poly-ADP-ribose and the acidic domain (AD) that is the histone chaperone domain. **B,** Sequence alignment of the C-terminal acidic domain of APLF, showing that it is highly conserved across tetrapod species, including the presence of aromatic residues within the acidic region (indicated with arrows) and a positively charged C-terminal region.

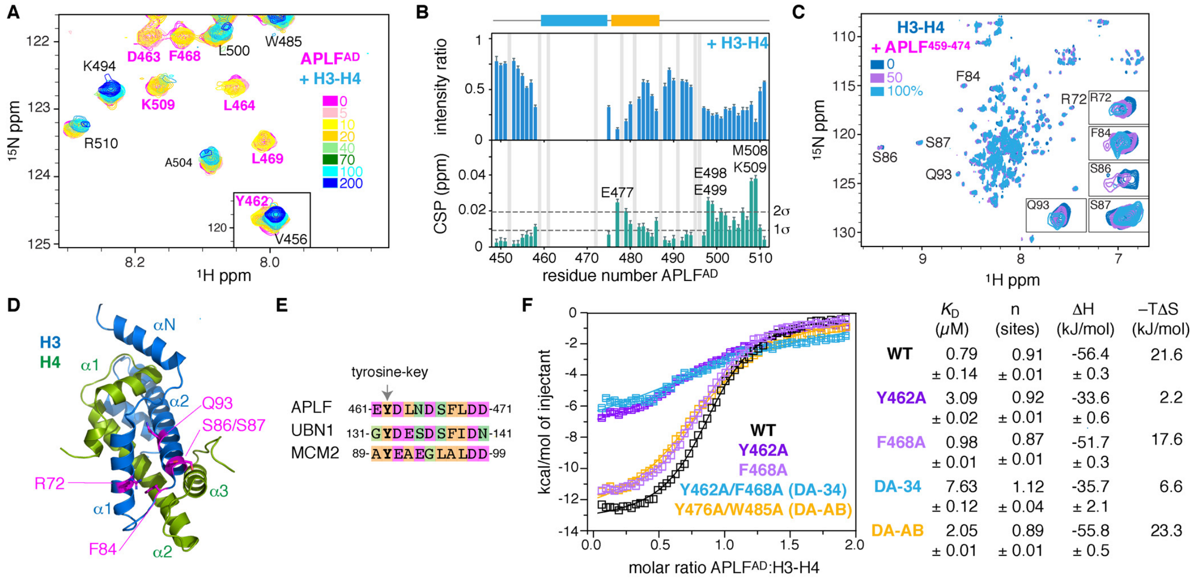


**Fig. S2 | APLF$^{AD}$ binds H3-H4 at the H3 α1-α2-patch through a conserved tyrosine. A**, Overlay of [$^{1}$H,$^{15}$N]-HSQC NMR spectra of APLF$^{AD}$ with increasing concentrations of H3-H4. Inset shows the V456/Y462 resonances. Color coding indicated in the Figure. Residues with disappeared resonances at the end of the titration are labeled magenta. **B**, Analysis of APLF$^{AD}$ peak intensity ratios (upper panel) and weighted chemical shift perturbations (CSPs) (lower panel) upon addition of 1 molar equivalent of H3-H4. Residues with CSPs larger than two standard deviations (σ) from the 10% trimmed mean (indicated by dashed line) are labeled. Residues without data (prolines/overlapped peaks) are in gray. **C**, Overlay of [$^{1}$H-$^{15}$N]-HSQC NMR spectra of H3-H4 with increasing concentrations of APLF peptide (res. 459-474) with insets showing the most affected peaks. Color coding indicated in the Figure. **D**, The H3 residues most affected by addition of APLF peptide cluster together in the α1-α2 region of H3 in the H3-H4 dimer structure. **E**, Sequence alignment of the H3-H4 interaction region in APLF with those of histone chaperones UBN1 and MCM2, highlighting the conservation of the tyrosine key (Y462). **F**, ITC binding isotherms and derived thermodynamic parameters, including best-fit values and fitting errors, of APLF$^{AD}$ (wild-type or indicated mutants) titrated to H3-H4, showing that mutation of Y462 results in a strong decrease of binding affinity and enthalpy.

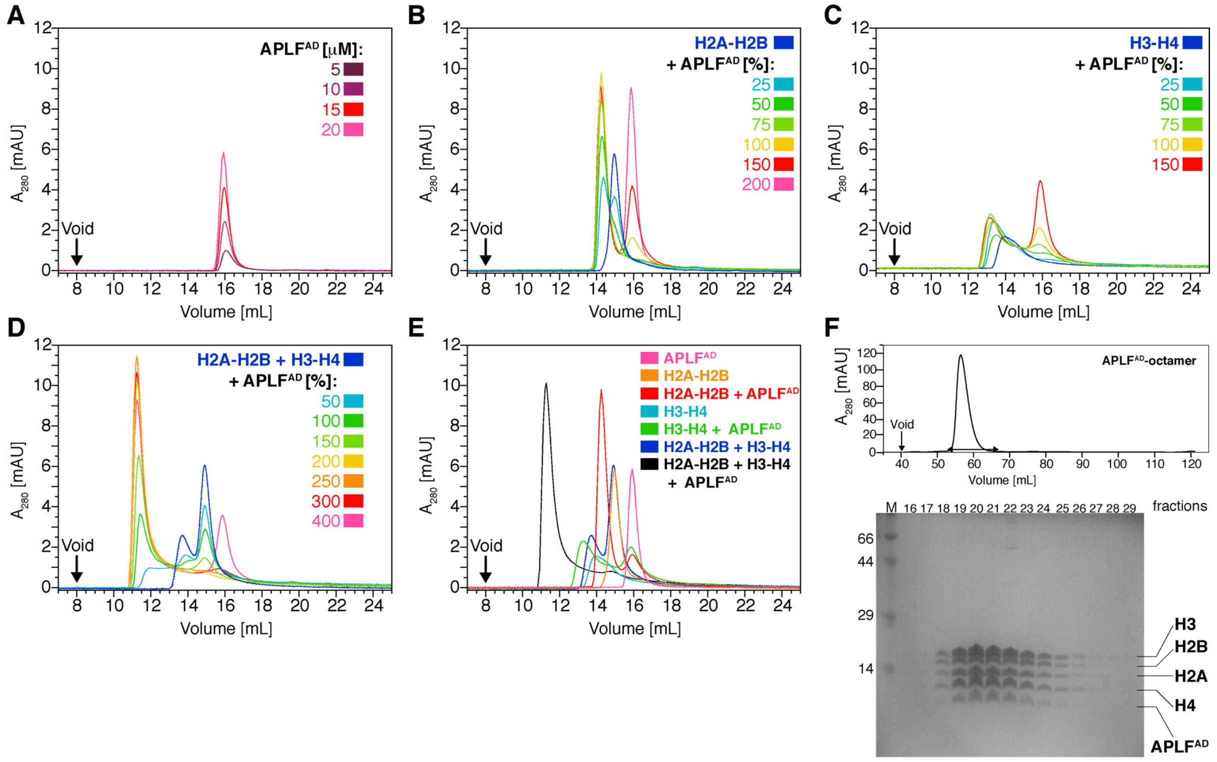


**Fig. S3 | Formation of histone–APLF$^{AD}$ complexes as shown from analytical size exclusion chromatography (SEC). A**, APLF$^{AD}$ alone at different concentrations does not form higher molecular weight species. **B**, H2A-H2B alone elutes as dimer and forms one specific higher molecular weight complex upon addition of APLF$^{AD}$. At more than 100% APLF$^{AD}$ (i.e., 1:1 ratio with H2A-H2B), unbound APLF$^{AD}$ is found. **C**, H3-H4 alone elutes in a broad peak due to an equilibrium between H3-H4 tetramers, i.e. $(H3\text{-}H4)_2$, and H3-H4 dimers, and forms one specific higher molecular weight complex upon addition of APLF$^{AD}$ until a ratio of 1:1. **D**, The stoichiometric mix of H2A-H2B and H3-H4 (octamer-mix) elutes in two peaks corresponding to H2A-H2B and H3-H4 dimers, and $(H3\text{-}H4)_2$ tetramers and forms one specific higher molecular weight complex upon addition of APLF$^{AD}$. The molar percentage of APLF$^{AD}$ added is expressed per molar equivalent of histone octamer, indicating a stoichiometry of the complex of 2 APLF$^{AD}$ per histone octamer. **E**, Overlay of chromatograms taken from **A-D** for direct comparison of the individual components and their complexes. **F**, Chromatogram (top) and SDS-PAGE analysis (bottom) of a gel filtration run of the octamer-mix with 2 equivalents of APLF$^{AD}$ added, showing that all five components are in the peak fractions (horizontal arrow in chromatogram) of the APLF$^{AD}$–H2A-H2B–H3-H4 complex.

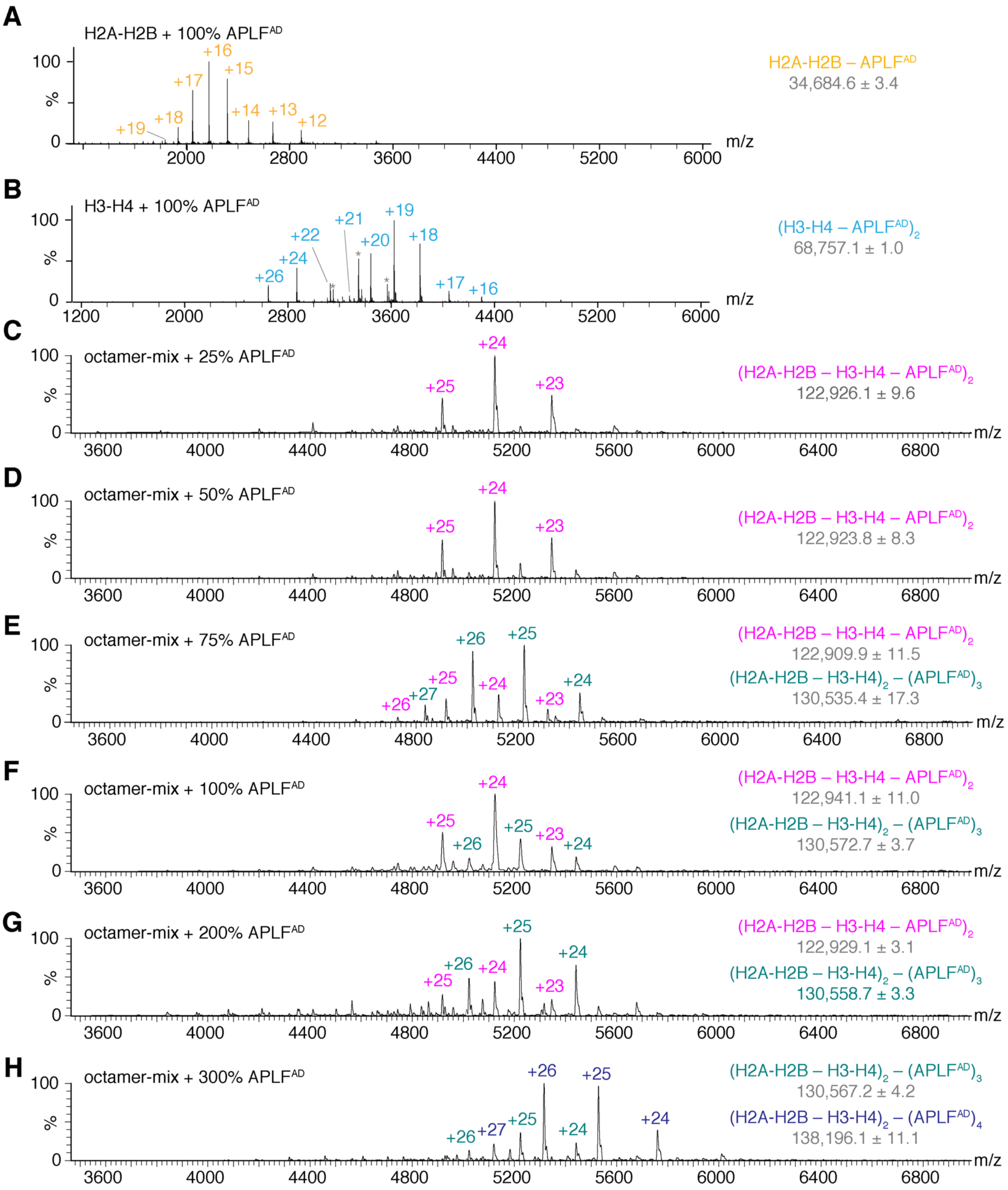


**Fig. S4 | Formation of a histone octamer–APLF$^{AD}$ complex as shown from native mass spectrometry analysis.** Native mass spectra of H2A-H2B (**A**), H3-H4 (**B**) and stoichiometric H2A-H2B and H3-H4 histone mix (octamer-mix, **C**-**H**) incubated with different concentrations of APLF$^{AD}$, expressed per histone dimer for H2A-H2B and H3-H4 and per octamer equivalent for the mix. **B**, Peaks labeled * in the mixture with H3-H4 correspond to a $(H3\text{-}H4)_2$ species. **C**-**H**, No octamer is detected in absence of APLF$^{AD}$. A complex corresponding to two APLF$^{AD}$ bound to a histone octamer is formed as the only detectable species at low molar ratios APLF$^{AD}$ added. At high APLF$^{AD}$ ratios, a 3$^{rd}$ and 4$^{th}$ molecule of APLF$^{AD}$ can associate with the histone octamer, likely due to stabilization of electrostatic interactions in the gas-phase, allowing detection of additional non-specifically bound APLF$^{AD}$. Theoretical masses of the histone-APLF$^{AD}$ complexes are 34,668.5 Da for the (APLF$^{AD}$–H2A-H2B) complex, 68,758.5 Da for the $(APLF^{AD}\text{–}H3\text{-}H4)_2$ complex 122,839.8 Da for the $(APLF^{AD}\text{–}H2A\text{-}H2B\text{–}H3\text{-}H4)_2$ complex, 130,467.7 Da for the $(APLF^{AD})_3$ – $(H2A\text{-}H2B\text{–}H3\text{-}H4)_2$ complex, and 138,095.5 Da for the $(APLF^{AD})_4$ – $(H2A\text{-}H2B\text{–}H3\text{-}H4)_2$ complex.

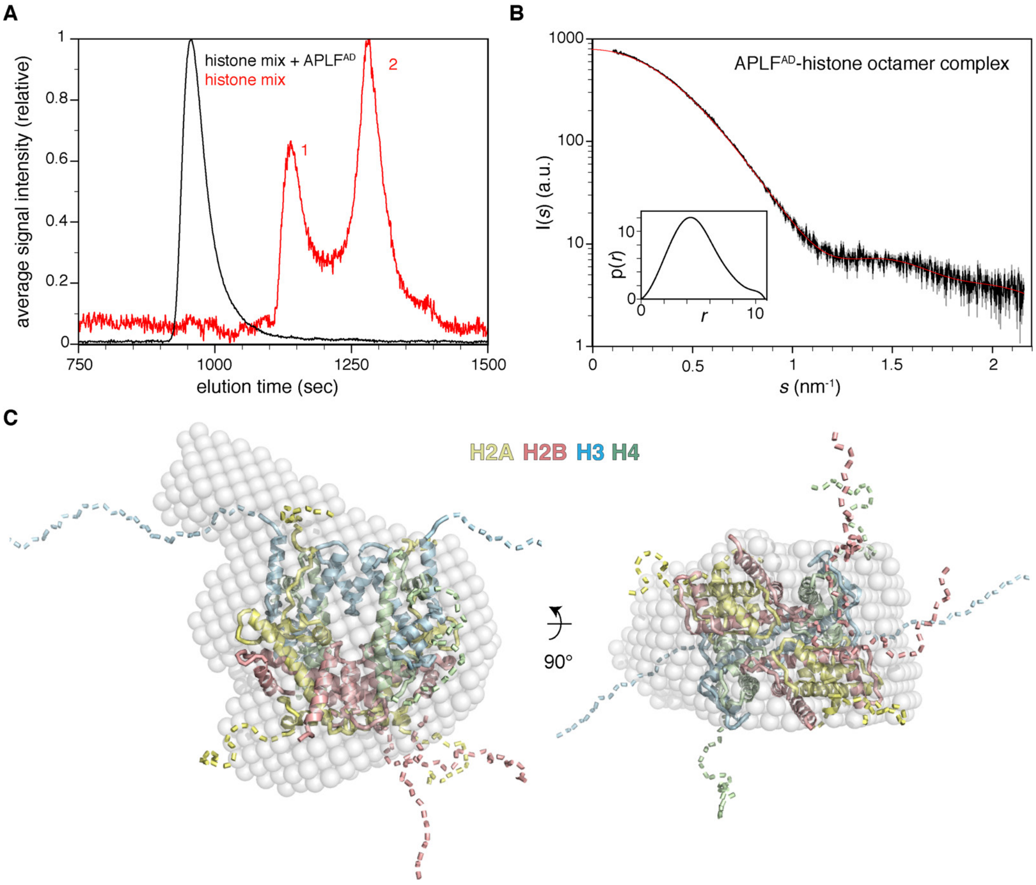


**Fig. S5 | SAXS analysis of the (APLF$^{AD}$–H2A-H2B–H3-H4)$_2$ complex shows that the complex dimensions are compatible with the histone octamer structure as in the nucleosome. A**, SEC-SAXS chromatograms of octamer-mix alone and in complex with APLF$^{AD}$. Two overlapping peaks are obtained for the octamer-mix with likely histone tetramers, (H3-H4)$_2$, contributing to peak 1 and histone dimers (H2A-H2B and H3-H4) to peak 2. A single complex is obtained in presence of APLF$^{AD}$, in agreement with the analytical SEC data (Fig. S3). Chromatograms were derived by integrating the intensity of the unsubtracted SAXS curves in each frame. **B**, Scattering curve of APLF$^{AD}$-histone octamer complex in solution. The inset shows the distance distribution. Analysis of the scattering curve (fit shown in red) showed that the APLF$^{AD}$-octamer complex has a radius of gyration ($R_g$) of 3.7 nm, a maximum particle dimension of 10.8 nm ($D_m$) and an estimated molecular weight (MW) of 130 kDa (111-142 kDa 95% confidence interval), in good agreement with expected MW for the APLF$^{AD}$-histone octamer complex (123 kDa). **C**, Superposition of the ab-initio SAXS-derived bead model with the histone octamer structure as in the nucleosome, showing that volume of the APLF$^{AD}$-octamer complex matches the dimensions of the histone octamer in the nucleosomal configuration. Flexible tails are schematically depicted as dashed lines.

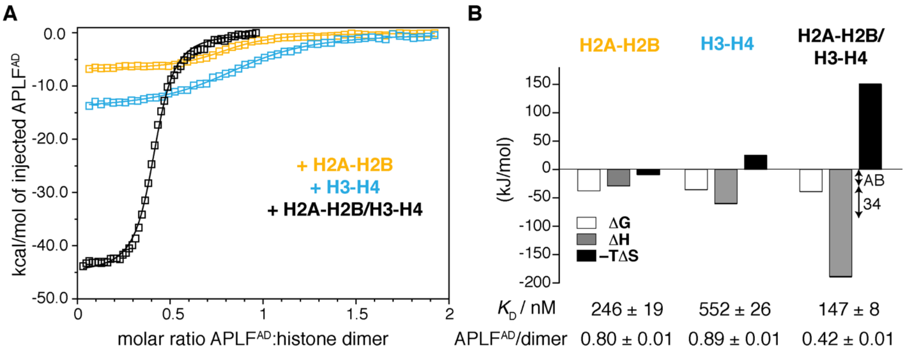


**Fig. S6 | ITC analysis of histone binding by APLF$^{AD}$ demonstrating the (APLF$^{AD}$–H2A-H2B–H3-H4)$_2$ complex is formed with high binding affinity and enthalpy. A,** ITC binding isotherms of APLF$^{AD}$ titrated to H2A-H2B, H3-H4 or octamer-mix, fitted to a one-set-of-sites binding model. The stoichiometry for octamer-mix binding is 0.5 APLF$^{AD}$ per histone dimer, i.e., one APLF$^{AD}$ binds two histone dimers, which is in line with the analytical gel filtration results (Fig. S3). The absence of a bi-phasic binding curve suggests that APLF$^{AD}$ binds H2A-H2B and H3-H4 in a concerted manner. **B**, Thermodynamic parameters for the APLF$^{AD}$-histone interaction, including best-fit values and fitting errors. APLF$^{AD}$ binds the histone octamer mix with a 1.5- and 3.8-fold higher affinity compared to the individual H2A-H2B or H3-H4 complexes. There is a remarkable increase in binding enthalpy (ΔH) for binding the octamer-mix compared to H2A-H2B and H3-H4 alone (see arrows in the Figure), indicating that APLF$^{AD}$ induces the formation of intermolecular contacts between H2A-H2B and H3-H4. This effect is accompanied by a strong entropic loss (ΔS) from the combination of at least three individual subunits forming a single complex. ΔG = binding free energy; ΔH = binding enthalpy, ΔS = binding entropy. Error bars on ΔG, ΔH and –TΔS are comparable to the line width of the bars.

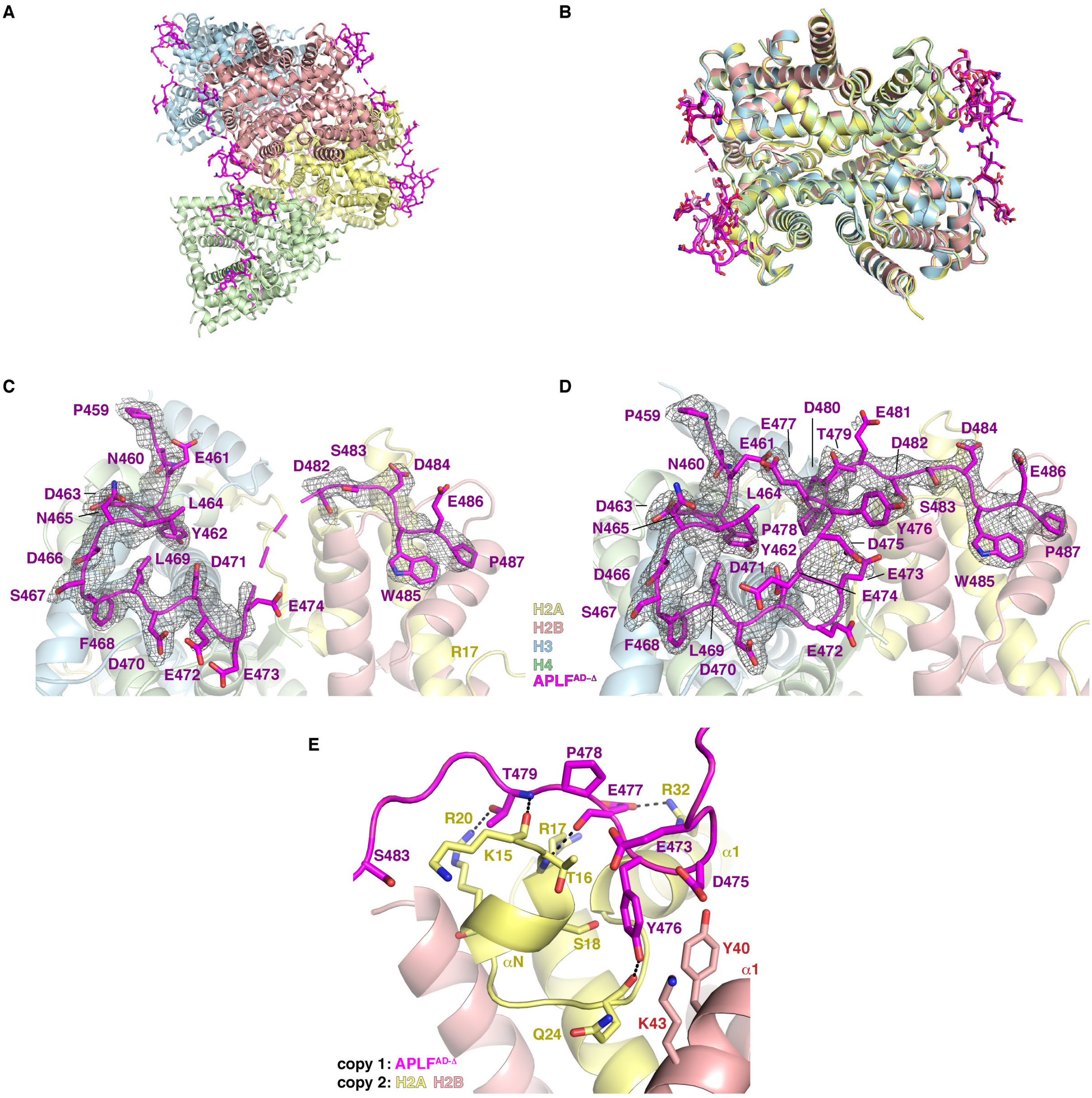


**Fig. S7 | Structure of the (APLF$^{AD\text{-}\Delta}$–H2A-H2B–H3-H4)$_2$ complex. A**, Ribbon view of the unit cell showing the four APLF$^{AD\text{-}\Delta}$-histone octamer complexes with APLF$^{AD\text{-}\Delta}$ (APLF residues 449-490) colored magenta in each complex and histone octamers colored differently in each complex. Each complex contains two copies of APLF$^{AD\text{-}\Delta}$, H2A-H2B and H3-H4. **B**, Overlay of the four complexes in the unit cell with APLF interface residues shown as sticks. Color coding as in panel **A**, with APLF$^{AD\text{-}\Delta}$ shown in different shades of purple. The overall average pairwise RMSD is 0.31 Å for all heavy backbone atoms. For APLF$^{AD\text{-}\Delta}$, the heavy atom average pairwise RMSD is 0.82 Å over all eight chains. **C**,**D** Electron density for one of the discontinuous (**C**) and the continuous (**D**) APLF$^{AD\text{-}\Delta}$ chains, contoured at one sigma. Selected residues are labeled. APLF$^{AD\text{-}\Delta}$ is in a coil conformation without any secondary structure, except for a turn conformation for residues L464-F468. **E**, Close view of the crystal packing interaction between two complexes involving the E472-E481 loop region of the continuous APLF$^{AD\text{-}\Delta}$ chain and the $\alpha$1-region of one H2A-H2B dimer in another histone octamer. Hydrogen bonds shown as dashes. Color coding indicated.

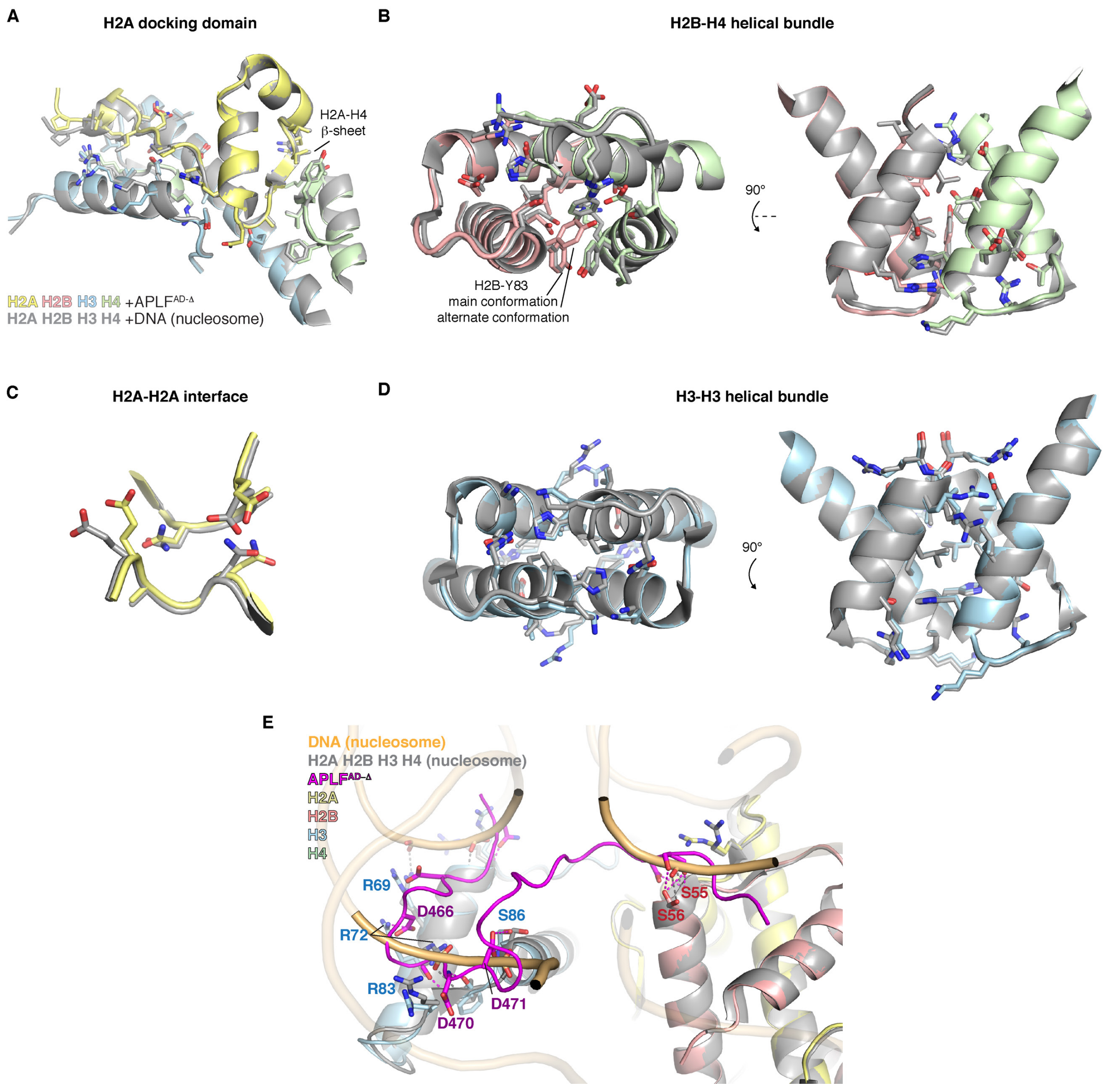


**Fig. S8 | Comparison of histone-histone contacts in the (APLF$^{AD-\Delta}$–H2A-H2B–H3-H4)$_2$ complex and in the nucleosome.** Structural superposition of the histone core structure in complex with residues 449-490 of APLF$^{AD}$ (APLF$^{AD-\Delta}$) or with DNA as part of the nucleosome (PDB 1KX5). **A**, The H2A docking domain interaction with H3-H4. The H2A-H4 β-sheet and the H3 αN-helix are indicated. Strikingly, these regions are not folded in the isolated histone complexes or when they are bound to other histone chaperones, and are not bound by APLF$^{AD-\Delta}$ directly. **B**, The H2B-H4 helical bundle. The main and alternate conformation of H2B Y83 in the APLF$^{AD-\Delta}$-complex are indicated. **C**, The H2A-H2A interface. **D**, Zoom on the H3-H3 helical bundle. Side chains of residues that have at least one interatomic distance < 4 Å are shown as sticks. Color coding indicated in the Figure. Formation of histone-histone contacts in the (APLF$^{AD-\Delta}$–H2A-H2B–H3-H4)$_2$ complex is likely promoted by tethering of H2A-H2B and H3-H4 by APLF$^{AD-\Delta}$ and further aided by stabilization of histone folding and reduction of the electrostatic repulsion. **E**, Zoom on the APLF$^{AD}$-histone binding interface with nucleosomal DNA superimposed (color coding indicated). Polar contacts indicated with dashed lines; selected residues are labeled. Several acidic residues of APLF$^{AD}$ form intermolecular hydrogen bonds with histone residues that are hydrogen bonded to DNA in the nucleosome.

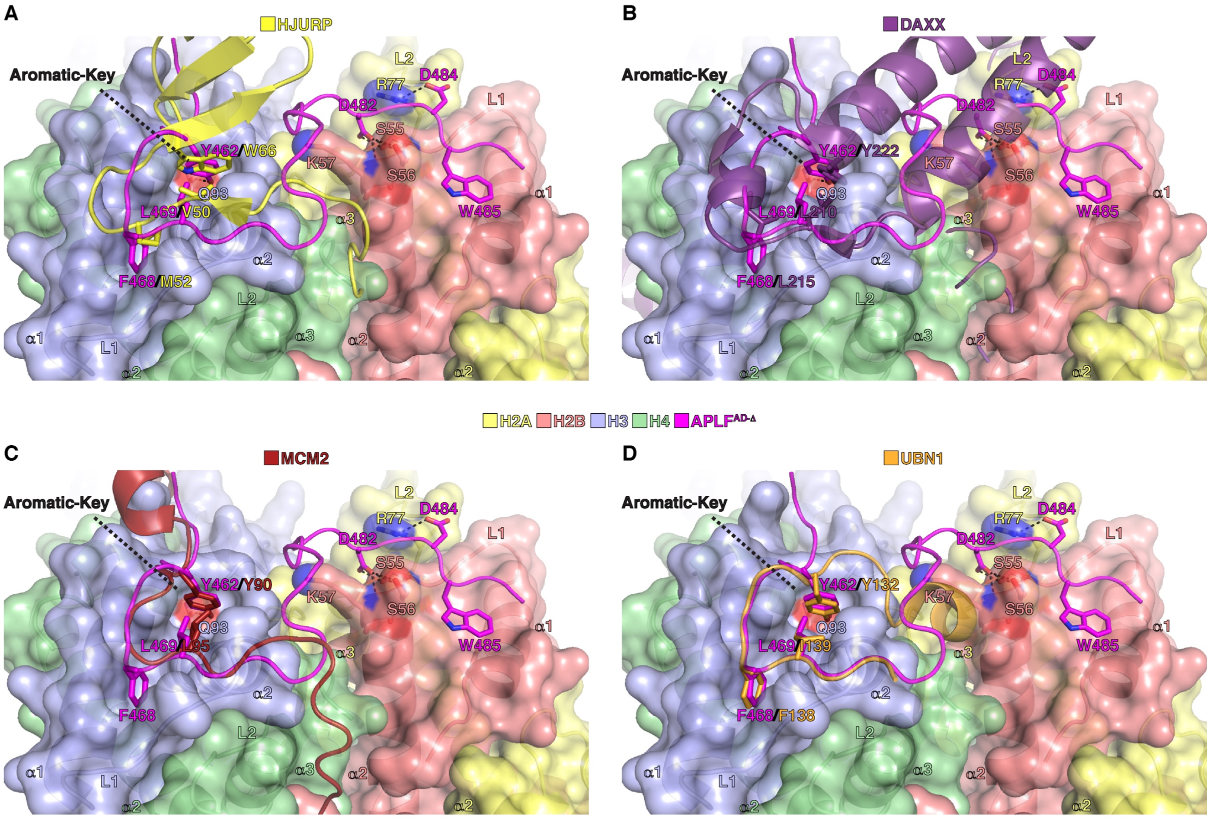


**Fig. S9| Structural comparison of different H3(.3)/CENP-A-H4 histone chaperones with APLF$^{AD}$. A**, HJURP (PDB: 3R45), **B**, DAXX (PDB: 4HGA), **C**, MCM2 (PDB: 5BNV), and **D**, UBN1 (PDB: 4ZBJ) in complex with H3(.3)/CENP-A-H4. All these chaperones feature an aromatic residue (HJURP W66, UBN1 Y132, MCM2 Y90, or DAXX Y222) that fits into a deep surface pocket on H3(.3)/CENP-A, where the aromatic ring forms extensive van der Waals interactions with histone residues, the aromatic-key motif. In UBN1, MCM2 and DAXX the tyrosine hydroxyl group forms an additional hydrogen bond with H3(.3) Q93.

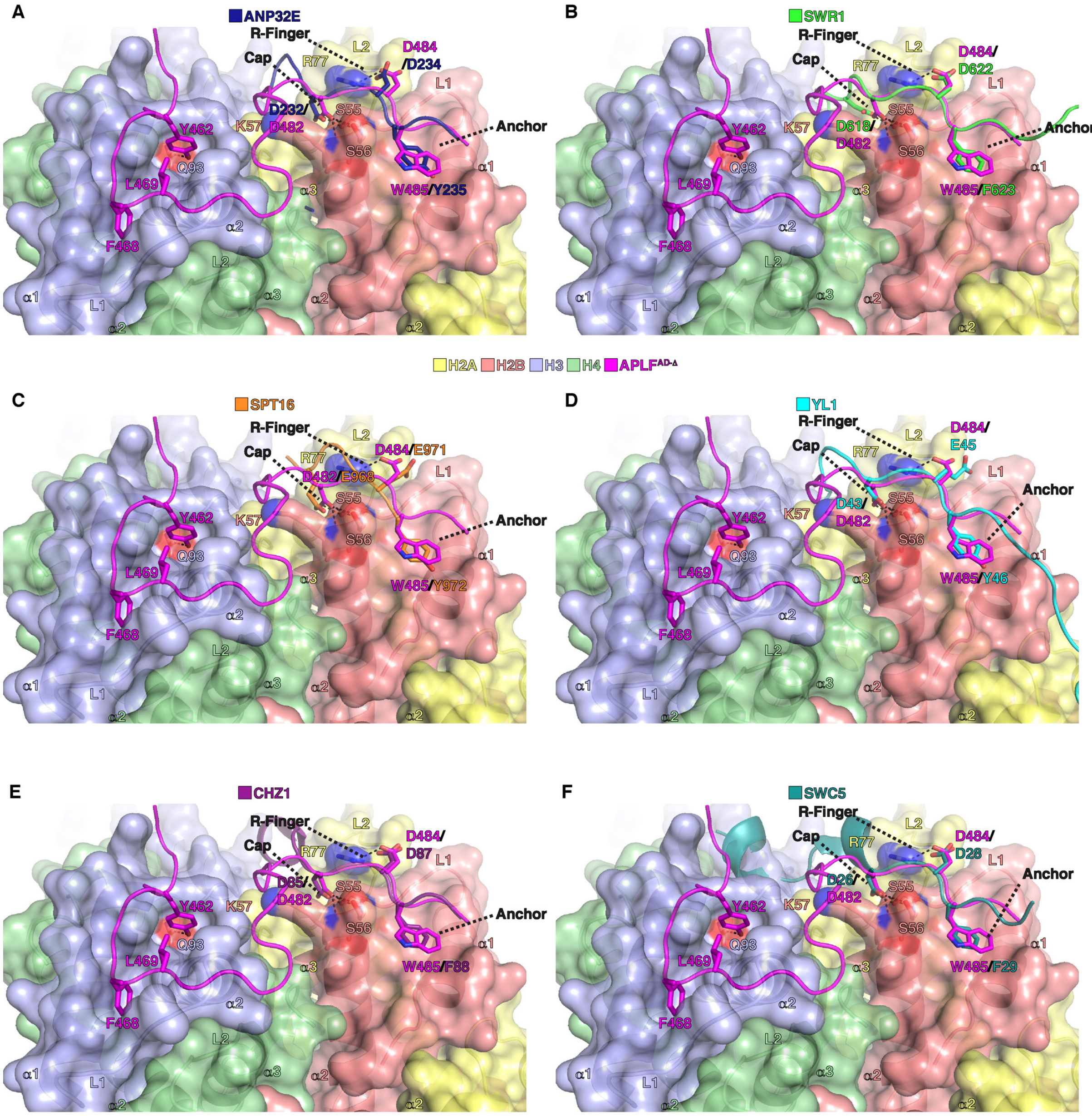


**Fig. S10 | Structural comparison of different H2A(.Z)-H2B histone chaperones with APLF$^{AD}$. A**, ANP32E (PDB: 4CAY), **B**, SWR1 (PDB: 4M6B), **C**, SPT16 (PDB: 4WNN), **D**, YL1 (PDB: 5CHL), **E**, CHZ1 (PDB: 6AE8), and **F**, SWC5 (PDB: 6KBB) in complex with H2A(.Z)-H2B. All these chaperones use an electrostatic R-finger binding motif, capping of the N-terminus of the H2B $\alpha$2-helix and an aromatic anchor (phenylalanine, tyrosine, or tryptophan) that fits into a surface pocket on H2A(.Z)-H2B where the aromatic ring forms van der Waals interactions with H2A(.Z)-H2B residues.

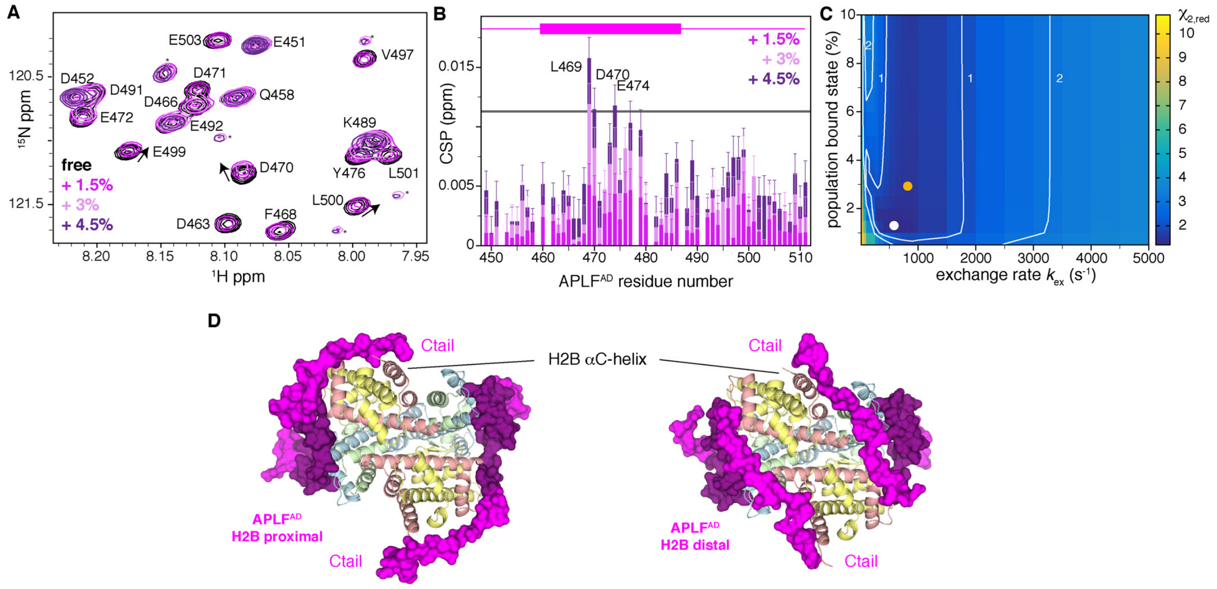


**Fig. S11 | NMR and XL-MS data indicate APLF$^{AD}$ envelops the histone octamer. A**, Overlay of [$^{1}$H,$^{15}$N]-TROSY NMR spectra of perdeuterated $^{15}$N-labeled APLF$^{AD}$ with increasing concentrations of octamer-mix. Color coding indicated in the Figure. Upon addition of half equivalent octamer mix (i.e. APLF$^{AD}$:octamer is 2:1) only signals for the N-terminal 8 residues L499-V456 could be detected, also when using a CRINEPT experiment. Unassigned signals indicated with *. **B**, Residues with largest chemical shift perturbations (CSPs) cluster in the region of the crystallized APLF fragment (shown as a purple box in top of the Figure). **C**, Error surface for a global two-state (free-bound) fit of the APLF$^{AD}$ relaxation dispersion data in presence of 1.5% octamer-mix (1:0.03:0.03 APLF$^{AD}$:H2A-H2B:H3-H4), showing the reduced $\chi^2$ as function of the exchange rate ($k_{ex}$) and the fractional population of the bound state ($p_B$). The best-fit combination ($p_B$ = 1.23 %; $k_{ex}$ = 530 $s^{-1}$) is indicated as white dot. The orange dot shows the best-fit combination when fixing the bound state population to 3% ($k_{ex}$ = 820 $s^{-1}$), as expected from the affinity measured by ITC (Fig. S6). **D**, The cross-links are compatible with APLF$^{AD}$ C-terminal region contacting either the proximal H2A-H2B dimer (bound at the H2B $\alpha$1-$\alpha$2-patch by the same APLF$^{AD}$ molecule) or the distal H2A-H2B dimer (bound at the H2B $\alpha$1-$\alpha$2-patch by the other APLF$^{AD}$ molecule). Positioned near the histone octamer surface of H2A-H2B, the C-terminal region will act as a steric block and limit DNA binding in either conformation.

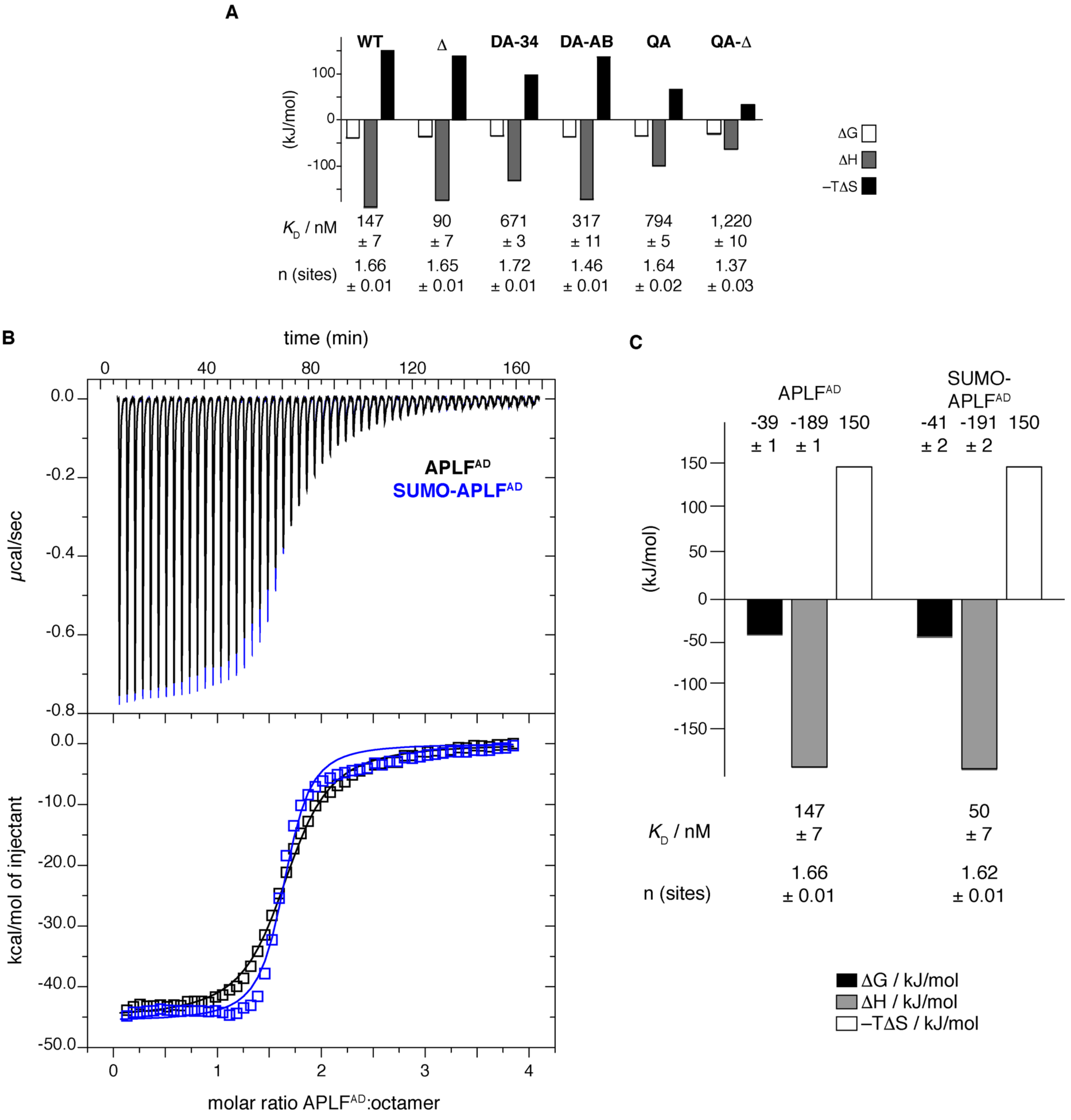


**Fig. S12 | Mutation of aromatic anchor residues in APLF$^{AD}$ reduced binding affinity and enthalpy. A,** ITC-derived thermodynamic parameters for the titrations of histone octamer-mix with APLF$^{AD}$ wild-type (WT), the deletion mutant used for crystallization (APLF$^{449\text{-}490}$, Δ), the double (DA) and quadruple (QA) aromatic anchor mutants (Y462A/F468A = DA-34; Y476A/W485A = DA-AB; Y462A/F468A/Y476A/W485A = QA), or the deletion mutant carrying the QA mutation (QA-Δ). Data for the QA-mutants was obtained using a SUMO-tagged version of the APLF$^{AD}$ protein. Data were fit to a one-set-of-sites binding model. The best-fit values and fitting errors are shown for the dissociation constant $K_D$ and the stoichiometry of binding $n$. **B**, ITC titration curves including raw data of APLF$^{AD}$ or SUMO-APLF$^{AD}$ titrated to the octamer mix, showing that the SUMO tag does not influence histone octamer-mix binding. The resulting binding isotherms were fit to a one-set-of-sites binding mode. **C**, ITC derived best-fit values and fitting errors are shown in the table, the derived thermodynamic parameters in the histogram. ΔG = binding free energy; ΔH = binding enthalpy, ΔS = binding entropy.

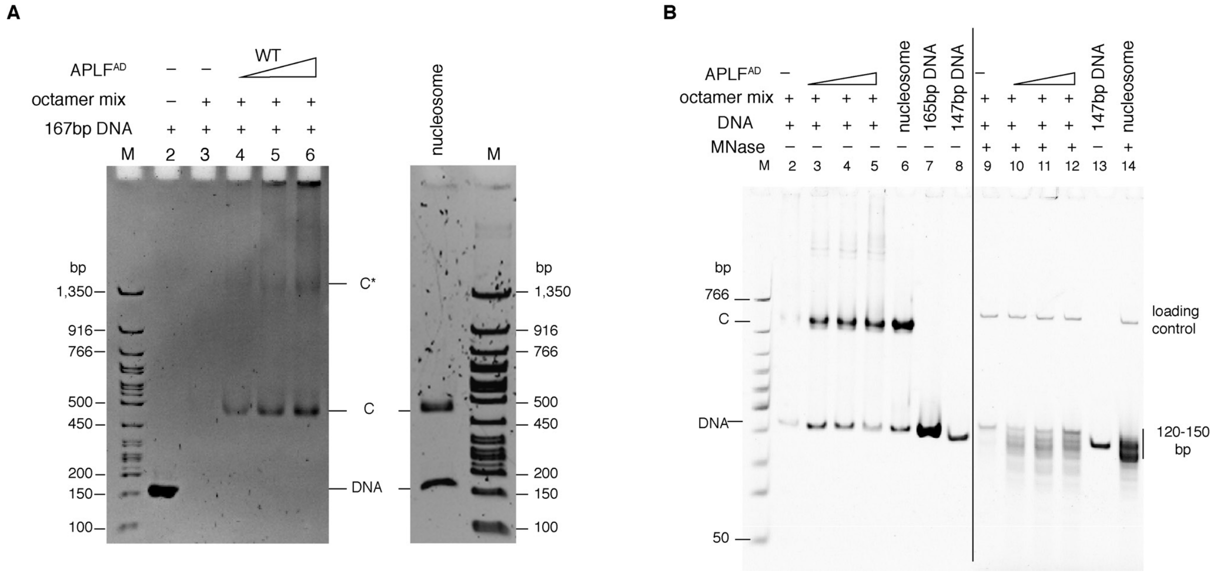


**Fig. S13 | APLF$^{AD}$ promotes nucleosome formation in vitro. A**, The electrophoretic mobility of the soluble protein-DNA complexes formed in presence of wild-type APLF$^{AD}$ during the precipitation-rescue assay (Main Text Fig. 3C; bands "C") corresponds to that of the nucleosome. **B**, Native PAGE analysis of soluble DNA (free or in histone complexes) before (lane 2-5) or after (lane 9-12) MNase digestion obtained after incubation of DNA with the octamer-mix with increasing concentrations of APLF$^{AD}$, together with controls. Octamer-mix (3 µM) is incubated directly with 165 bp DNA (1 µM) (lane 2) or first pre-incubated with 50, 100, or 200 µM wild-type (WT) APLF$^{AD}$ (lane 3–5), after which the soluble protein-DNA complexes are analyzed by native PAGE. Salt-gradient deposited nucleosome (lane 6) and free DNA (lane 7) are shown as controls. The free DNA band and the band corresponding to nucleosome (C) are indicated. Compared to octamer-mix and DNA alone, addition of APLF$^{AD}$ results in a clear increase in intensity for the nucleosomal band (lane 3–5 vs. 2). To assess the degree of DNA protection, the DNA in the soluble complexes was digested by micrococcal nuclease (MNase) and purified (lane 9-12). Compared to octamer-mix and DNA alone (lane 9), addition of APLF$^{AD}$ (lane 10–12) results in a clear increase in intensity in the range 120–150 bp. Nucleosomes give rise to protection of 120-150 bp DNA from MNase digestion. Fragments obtained after MNase digestion of salt-gradient deposited nucleosome are shown in lane 14 with undigested 147 bp free DNA as control in lane 7 and 13; DNA marker in lane M. Samples after MNase treatment contain a loading control DNA fragment (indicated with *). Gels of three repeat experiments are available in source data. Quantification of the signal intensity of bands in the three repeats corresponding to DNA fragments smaller than 127 bp, between 127 and 156 bp, and larger than 156 bp is shown in the Main Text.

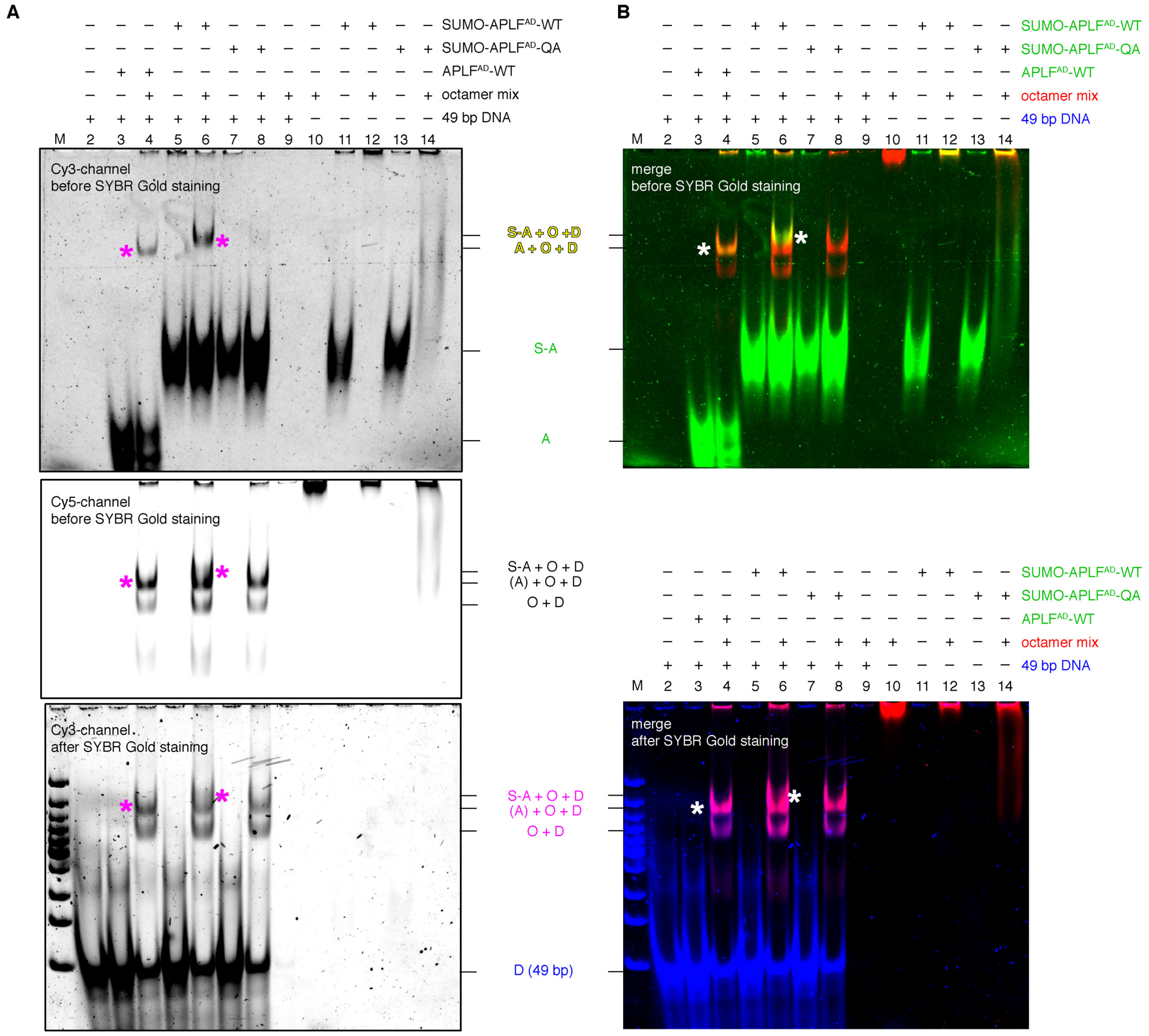


**Fig. S14 | The (APLF$^{AD}$–H2A-H2B–H3-H4)$_2$ complex can bind short double-stranded DNA to form a ternary complex.** Native PAGE analysis of indicated mixtures of TAMRA-tagged APLF$^{AD}$ proteins, histone octamer with Alexa647-labeled H2B, and DNA. Samples were crosslinked with DSS before separation on gel and visualization of APLF$^{AD}$ (green), H2B (red) and SYBR-Gold-stained DNA (blue) by fluorescence. **A**, Individual gel images of APLF$^{AD}$ (top), histone octamer (middle) and DNA (bottom) of the data shown in Main Text Fig. 4D,E. **B**, Merge of the gel images in panel **A** obtained before (top) and after (bottom) SYBR gold DNA staining. A = APLF$^{-AD}$, S-A = SUMO-APLF$^{AD}$, O = histone octamer, D = DNA. Labels are color-coded according to the fluorescent dye used and the expected merged color for each complex. Bands of the ternary complex are marked (*).

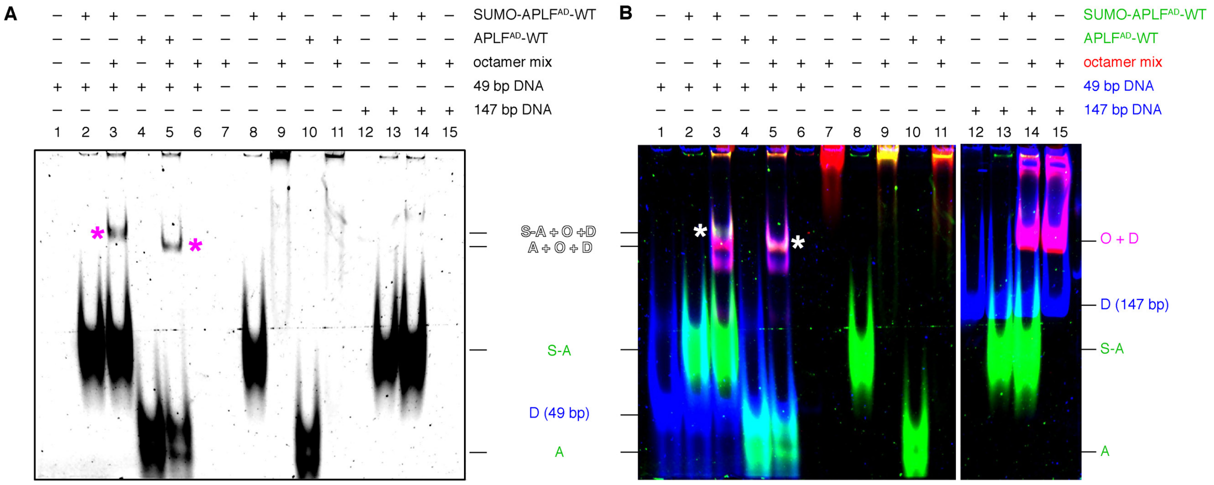


**Fig. S15 | The ternary APLF$^{AD}$-octamer-DNA complex forms on 49 bp but not on 147 bp DNA**. Native PAGE analysis of indicated mixtures of TAMRA-tagged APLF$^{AD}$ proteins, histone octamer with Alexa647-labeled H2B, and DNA. Samples were crosslinked with DSS before separation on gel and visualization of APLF$^{AD}$ (green), H2B (red) and SYBR-Gold-stained DNA (blue) by fluorescence. **A,** Cy3-scan with APLF$^{AD}$ signal before DNA staining. **B**, Merged image of the APLF$^{AD}$, histone and DNA scans. Due to the different DNA sizes, different scaling of each channel was applied for the 49 bp and 147 bp parts of the gel. Presence of APLF$^{AD}$ results in the formation of a ternary complex (marked bands) for 49 bp DNA (lane 3 and 5), but not for 147 bp DNA, where only binary DNA-octamer complexes are formed (magenta bands in **B**, lane 14). A = APLF$^{-AD}$, S-A = SUMO-APLF$^{AD}$, O = histone octamer, D = DNA. Labels are color-coded according to the fluorescent dye used and the expected merged color for each complex.

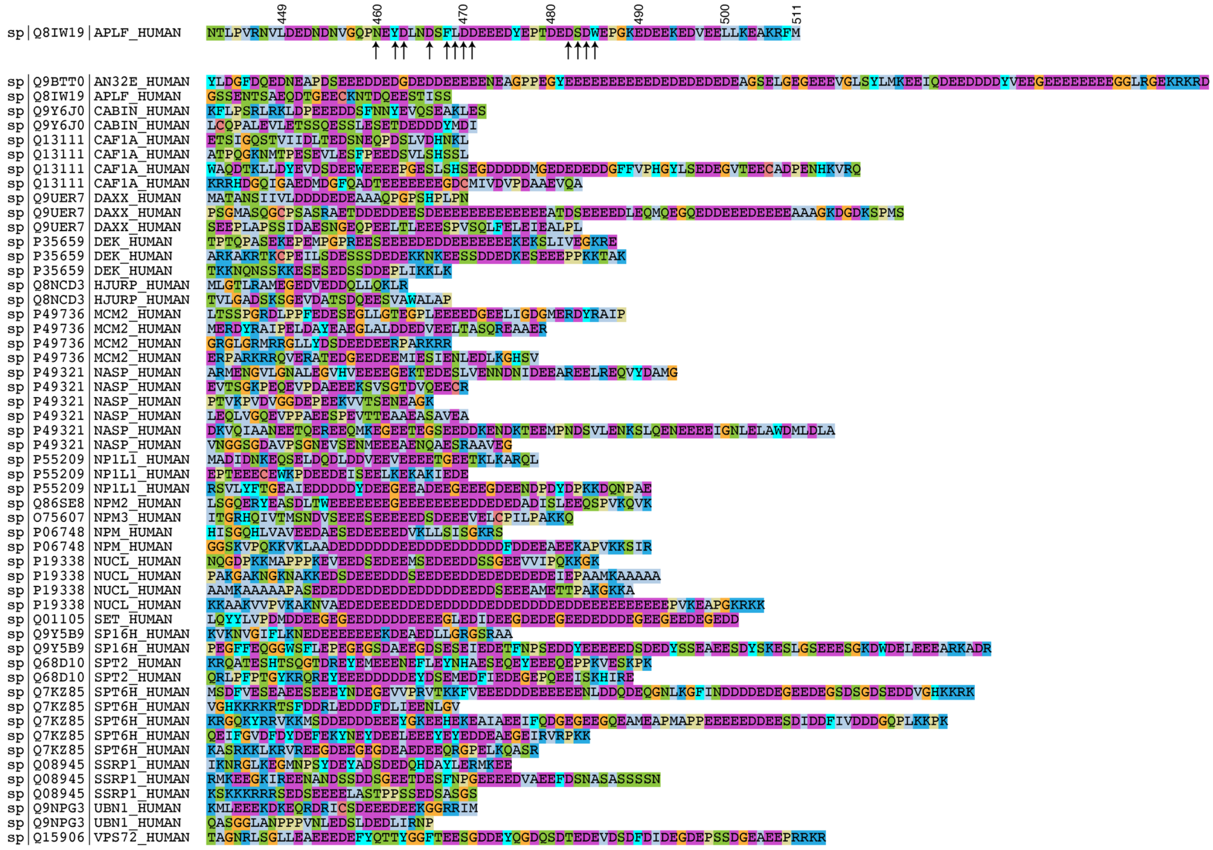

**Fig. S16 | Acidic domain sequences in human histone chaperones.** Acidic stretches in a list of 25 human chaperones(*32*) were identified as stretches of at least 15 residues with average local charge of –0.3 or lower and average disorder probability score of 0.6. Local charge was calculated by the EMBOSS server and disorder probability using the DISOPRED3 server. Amino acids are color coded by physiochemical property. All extracted acidic domain sequences, including that of APLF$^{AD}$ shown on top, are shown with 5 flanking residues on each end, and annotated with their Uniprot accession code. Several chaperones contain more than one acidic stretch that fits the selection criteria. Residues involved in intermolecular interactions with the histone octamer surface in the APLF$^{AD\text{-}\Delta}$–histone octamer crystal structure are indicated using arrows.

**Supplementary Table S1 | Crystallography data collection and refinement statistics**

| | |
|---|---|
| **Data collection** | |
| Beam line | X6A |
| Detector | Pilatus 6M-F |
| Space group | P212121 |
| Unit cell parameters: | |
| a, b, c (Å) | 105.2, 189.7, 204.3 |
| α, β, γ | 90, 90, 90 |
| copies per asymmetric unit | 4 |
| Matthew’s coefficient ($Å^3$/Da) | 2.58 |
| Resolution range (Å) | 20-2.35 (2.49-2.35)[a] |
| Wavelength | 1.0 |
| Unique reflections | 206799 (23719)[a] |
| Completeness (%) | 99.7 (99.7)[a] |
| Redundancy | 6.7 (6.7)[a] |
| $R_{meas}$(%) | 12.5 (196.6)[a] |
| I/ σ(I) | 14.41 (1.03)[a] |
| Wilson B-factor ($Å^2$) | 49.0 |
| CC ½ (%) | 99.9 (42.0)[a] |
| **Refinement** | |
| Resolution range | 20-2.35 |
| Reflection (test set) | 169778 (8284)[a] |
| Total number of atoms | 24906 |
| Protein atoms | 24400 |
| Water and solvent atoms | 427 |
| Other atoms | 79 |
| $R_{work}$/$R_{free}$ (%) | 17.6/23.4 |
| Average B-factor (overall) | 27.4 |
| Average B-factor (protein) | 27.4 |
| Average B-factor (water) | 38.0 |
| RMSD from ideal bond length | 0.008 |
| RMSD from ideal bond angle | 0.926 |
| Ramachandran plot: | |
| Preferred (%) | 97.87 |
| Allowed (%) | 99.76 |
| Outliers (%) | 0.24 |

[a] data in high-resolution shell

**Supplementary Table S2 | SAXS data collection and analysis**

| Histone H2A:H2B:H3:H4 complex with APLF$^{AD}$ | |
|---|---|
| Source | *Homo sapiens(APLF$^{AD}$),*<br>*Drosophila melanogaster(H2A, H2B, H3, H4)* |
| SEC-SAXS buffer | 25 mM NaPi, 300 mM NaCl, 3% v/v glycerol, 1 mM DTT, pH 7 |
| SEC-SAXS column | Superdex 200 inc. 10/300 |
| SEC-SAXS injection volume | 40 µL |
| SEC-SAXS injection concentration | 14 mg/mL |
| UniProt entry<br>(amino acid sequence range) | APLF$^{AD}$ (Q8IW19 (449-511), His2A (P84051 (2-124)), His2B (P02283 (2-123), His3 (P02299 (2-136)), His4 P84040 (2-103) |
| Wavelength (nm) | 0.12 |
| q range ($nm^{-1}$) | 0.03–5.0 |
| Temperature (K) | 293 |
| Exposure time (s) | 40 |
| **Structural Parameters** | |
| $R_G$ (nm), Guinier | 3.7 ± 0.1 |
| $R_G$ (nm), P(r) | 3.7 ± 0.1 |
| $D_{max}$ (nm) | 10.8 ± 0.5 |
| **Molecular Weight Determination** | |
| $MW_{exp}$ (kDa) | 122 |
| $MW_{QP}$ (kDa) | 134 ± 10 |
| $MW_{MoW}$ (kDa) | 94 ± 10 |
| $MW_{VC}$ (kDa) | 120 ± 10 |
| $MW_{S\&S}$ (kDa) | 154 ± 10 |
| $MW_{Bayesian}$ (kDa) | 130.9 |

| | |
|---|---|
| Cred. interval, (kDa) | 111.3–142.2 |
| range (%) | 95.68 |
| **Software Employed** | |
| Primary data reduction | SASFLOW/CHROMIXS |
| Data processing | PRIMUS |
| *Ab initio* modeling | DAMMIF |
| SASBDB ID | SASDJJ5 |

**Supplementary Table S3 | List of APLF mutants used in the study**

| Short name (long name) | Mutation |
|---|---|
| DA-AB<br>(double anchor mutant for H2A/H2B interface) | Y476A, W485A |
| DA-34<br>(double anchor mutant for H3/H4 interface) | Y462A, F468A |
| QA<br>(quadruple anchor mutant) | Y462A, F468A, Y476A, W485A |
| Δ<br>(truncation mutant) | D491STOP |
| QA-Δ<br>(truncated quadruple anchor mutant) | Y462A, F468A, Y476A, W485A, D491STOP |
| ΔAD<br>(full-length APLF w/ deletion of acidic domain) | D450STOP |

**Supplementary Table S4 | List of primers used to generate the APLF$^{AD}$ transfection vectors**

| Name | Sequence (5’-3’) |
|---|---|
| XhoI FW | TTAACATCTCGAGCTCTGTCCGGGGGCTTCGAGCTG |
| APLF RV | CGAAGCTTGGGTACCACGCGTGAATTCGGATTCCTA |
| APLF Y462A/F468 FW | CAACCCAATGAGGCTGACCTGAACGACAGCGCGCTAGATGATGAG |
| APLF Y462A/F468 RV | GAGTAGTAGATCGCGCGACAGCAAGTCCAGTCGGAGTAACCCAAC |
| APLF Y476A/W485A FW | GAAGACGCTGAGCCAACAGATGAAGATTCTGACGCTGAACCAGGAAAGG |
| APLF Y476A/W485A RV | CCTTTCCTGGTTCAGCGTCAGAATCTTCATCTGTTGGCTCAGCGTCTTC |
| APLF ∆AD (D450STOP) FW | CTTCCAGTGAGAAATGTTTTA-Δ-TAGGAATCCGAATTCACGCG |
| APLF ∆AD (D450STOP) FW | CCATGGTGCGCACTTAAGCCTAAGGAT-Δ-ATTTTGTAAAGAGTG |

**Supplementary Table S5. List of primary antibodies**

| Protein | Host | Source | Western blot |
|---|---|---|---|
| APLF | Rabbit | Gift from A. Yasui | 1:500 |
| H4 | Mouse | Upstate #07-108 | 1:1500 |
| H2B | Rabbit | Millipore #07-371 | 1:1500 |
